# RAD sequencing enables unprecedented phylogenetic resolution and objective species delimitation in recalcitrant divergent taxa

**DOI:** 10.1101/019745

**Authors:** Santiago Herrera, Timothy M. Shank

## Abstract

Species delimitation is problematic in many taxa due to the difficulty of evaluating predictions from species delimitation hypotheses, which chiefly relay on subjective interpretations of morphological observations and/or DNA sequence data. This problem is exacerbated in recalcitrant taxa for which genetic resources are scarce and inadequate to resolve questions regarding evolutionary relationships and uniqueness. In this case study we demonstrate the empirical utility of restriction site associated DNA sequencing (RAD-seq) by unambiguously resolving phylogenetic relationships among recalcitrant octocoral taxa with divergences greater than 80 million years. We objectively infer robust species boundaries in the genus *Paragorgia,* which contains some of the most important ecosystem engineers in the deep-sea, by testing alternative taxonomy-guided or unguided species delimitation hypotheses using the Bayes factors delimitation method (BFD*) with genome-wide single nucleotide polymorphism data. We present conclusive evidence rejecting the current morphological species delimitation model for the genus *Paragorgia* and indicating the presence of cryptic species boundaries associated with environmental variables. We argue that the suitability limits of RAD-seq for phylogenetic inferences in divergent taxa cannot be assessed in terms of absolute time, but depend on taxon-specific factors such as mutation rate, generation time and effective population size. We show that classic morphological taxonomy can greatly benefit from integrative approaches that provide objective tests to species delimitation hypothesis. Our results pave the way for addressing further questions in biogeography, species ranges, community ecology, population dynamics, conservation, and evolution in octocorals and other marine taxa.

## INTRODUCTION

Species delimitation is problematic in many taxa due to the difficulty of evaluating predictions from species delimitation hypotheses derived using different species concepts. Species concepts set particular expectations of the properties used to support species delimitations (De Queiroz 2007). For example, the classic biological species concept requires intrinsic reproductive isolation between heterospecific organisms and interbreeding among homospecific organisms resulting in viable and fertile descendants (Mayr 1942; Dobzhansky 1970). In many cases, if not the majority, it is difficult to evaluate behavioral, reproductive, and ecological properties due to technical limitations of field or laboratory work, which largely determine the kind of observations and data that can be obtained. In these cases researchers conventionally rely on morphological observations and/or DNA sequence data to generate species delimitation hypotheses.

Although there have been significant attempts at developing statistical methods to objectively identify species-diagnostic morphological discontinuities (e.g., Zapata & Jimenez 2012), most species delimitations continue to be performed subjectively based on assessments made by specialized taxonomists. Molecular phylogenetic analyses of DNA sequences provide an independent way to test these species delimitation hypotheses utilizing a variety of methods, ranging from variability thresholds of barcode sequences (Hebert *et al.* 2003), to probabilistic coalescent-based model methods (Pons *et al.* 2006; Yang & Rannala 2010; Fujisawa & Barraclough 2013; Grummer *et al.* 2014). These molecular methods rely on informative DNA sequence markers, and in many cases on resolved phylogenies.

The sub-class Octocorallia (Phylum Cnidaria), which includes animals known as gorgonians, sea pens, and soft corals, is an example of a recalcitrant group where species delimitations are problematic. Octocorals are predominantly a deep-sea group (Cairns 2007; Roberts & Cairns 2014) and therefore are extremely difficult to observe and collect. Classic morphology-based species delimitation and identification in this group is arduous for non-specialists, and challenging to replicate among taxonomists (Daly *et al.* 2007; McFadden *et al.* 2010b). Variations in octocoral colony architecture and micro-skeletal structures – sclerites – are used as species diagnostic characters (Bayer 1956). However, studies over the last 15 years have shown that in many cases species delimitations and systematics based on these morphological traits keep little to no correspondence with the patterns of genetic diversity and relatedness inferred using mitochondrial and ribosomal DNA sequence markers (McFadden *et al.* 2006; Clark *et al.* 2007; France 2007; Dueñas & Sánchez 2009). A confounding factor when analyzing mitochondrial DNA markers is the fact that anthozoans, including octocorals, have slow rates of sequence evolution relative to other metazoans (Shearer *et al.* 2002; Hellberg 2006). Furthermore, octocoral mitochondrion is unique among eukaryotes by having a functional DNA mismatch repair gene — *mtMutS —* which presumably is responsible for the extremely low sequence variability observed in this group (Bilewitch & Degnan 2011). Traditional molecular markers have thus been remarkably insufficient to resolve relationships at all taxonomic levels within the octocorals (Berntson *et al.* 2001; France *et al.* 2002; Mcfadden *et al.* 2004; Smith *et al.* 2004; Thoma *et al.* 2009; Dueñas *et al.* 2014). Alternative nuclear markers, such as the ITS2 and *SRP54* have been used to examine interspecific and intraspecific relationships (Aguilar & Sánchez 2007; Concepcion *et al.* 2007; Grajales *et al.* 2007; Herrera *et al.* 2010); however, their application and impact has been limited due to issues regarding intragenomic variability (Sanchez & Dorado 2008) and low sequencing reliability (Mcfadden *et al.* 2010a). These long-standing technical problems have caused fundamental questions in octocorals regarding species differentiation, systematics, diversity, biogeography, and species ranges to remain unanswered.

Technological developments in next-generation sequencing platforms and library preparation methodologies have made genomic resources increasingly accessible and available for the study of non-model organisms, thus offering a great opportunity to overcome the difficulties inherent to the use of traditional sequencing approaches. One of these methodologies is restriction-site-associated DNA sequencing (RAD-seq), which combines enzymatic fragmentation of genomic DNA with high-throughput sequencing for the generation of large numbers of markers (Baird *et al.* 2008). RAD-seq has shown great promise to resolve difficult phylogenetic, phylogeographic, and species delimitation questions in diverse groups of eukaryotes (Emerson *et al.* 2010; Nadeau *et al.* 2012; Wagner *et al.* 2012; Eaton & Ree 2013; Jones *et al.* 2013; Cruaud *et al.* 2014; Escudero *et al.* 2014; Hipp *et al.* 2014; Leache *et al.* 2014; Herrera *et al.* 2015), including cnidarians (Reitzel *et al.* 2013) and most recently deep-sea octocorals (Pante *et al.* 2014). The number of orthologous restriction sites that can be retained across taxa, which decreases as divergence increases, limits the usefulness of RAD-seq for these kinds of studies. *In silico* studies in model organisms indicate that RAD-seq can be used to infer phylogenetic relationships in young groups of species (up to 60 million years old), such as *Drosophila* (Rubin *et al.* 2012; Cariou *et al.* 2013; Seetharam & Stuart 2013); however, the real limits of this technique have not been significantly explored.

In this study we aim to empirically explore the limits of RAD-seq to solve questions in phylogenetics and species delimitation. We focus on the recalcitrant *Anthomastus-Corallium* clade of octocorals (sensu McFadden *et al.* 2006) to test the utility of RAD-seq to resolve phylogenetic relationships among divergent taxa, and to infer objective species boundaries. Corals in the *Anthomastus-Corallium* clade (hereafter referred as the AC clade) are among the most conspicuous, widely distributed, and ecologically important benthic invertebrates in deep-water ecosystems (Roberts *et al.* 2009; Wating *et al.* 2011). This clade is constituted by more than 100 species defined morphologically, divided in 10 genera, and three families (World Register of Marine Species at http://www.marinespecies.org accessed on 2014-10-10), spanning a divergence time of over 100 million years (Ardila *et al.* 2012; Herrera *et al.* 2012). However, species delimitations and phylogenetic relationships in this clade, as in other octocorals, are controversial and conflictive (Herrera *et al.* 2010; Ardila *et al.* 2012; Herrera *et al.* 2012). Many of the species in this group are considered species indicators of Vulnerable Marine Ecosystems (e.g. ICES 2013), with some of them considered endangered (CITES 2014). Accurate species identifications, as well as complete inventories and knowledge of species ranges, are therefore critical to ensure the effectiveness and appropriateness of conservation and management policies.

## RESULTS

### Morphological species identifications

Using current species descriptions, colony observations, and scanning electron microscopy of sclerites, we identified a total of 12 putative morphological species among the 44 examined specimens from the AC clade (Table S1, Supplementary Material images). These species correspond to the genera *Paragorgia* (*P. arborea, P. stephencairnsi, P. johnsoni, P. maunga, P. alisonae, P. kaupeka,* and *P. coralloides*) and *Sibogagorgia* (*S. cauliflora*) of the family Paragorgiidae; *Hemicorallium* (*H. laauense-imperiale*) and *Corallium* of the family Coralliidae; and *Anthomastus* and *Heteropolypus* of the family Alcyoniidae.

### Octocorals are amenable to RAD sequencing

We generated a dense genome-wide set of genetic markers from the 44 AC clade specimens via RAD sequencing, using the 6-cutter restriction enzyme PstI, and used them to perform phylogenetic inferences and species delimitation analyses. We obtained roughly 3.9 ± 1.4 million reads (average ± standard deviation) per individual, of which 74.3 ± 8.1% were retained after stringent quality filtering steps (Table S2).

### Optimization of RAD-loci clustering parameters

To examine the sensitivity of the phylogenetic inference to the clustering parameters used to identify loci and create nucleotide matrices in the program pyRAD (Eaton 2014), we investigated different combinations of clustering thresholds (c 0.80, 0.85 and 0.90) and minimum number of taxa per locus (m 4, 6, and 9) in a reduced **‘backbone’** matrix (hereafter matrix names will be highlighted in bold) containing one individual from each of the 12 morphological species. The 9 resulting **backbone** matrices ranged in the total number of loci per matrix from approximately 9 to 60 thousand loci, increasing dramatically as the minimum number of taxa per locus was reduced (Table S3). In contrast, the different clustering thresholds did not have a significant effect on the total number of loci, but rather on the number of variable sites and, most importantly, on the number of phylogenetically informative sites (Table S3). Each resulting **backbone** matrix analyzed in RAxML (Stamatakis 2006) produced identical strongly-supported tree topologies (Fig S1). We selected c 0.80 (80% similarity among sequences) and m 9 (minimum coverage of taxa per locus of 75%) as the optimal combination of loci-clustering parameters because they minimized the proportion of missing data (0.20) in the matrix while maximizing the fraction of variable sites that were phylogenetically informative (0.24) (Table S3). The proportion of shared loci among individuals of Paragorgiidae and Coralliidae, lineages whose split has been estimated to be between 80-150 million years ago (Ardila *et al.* 2012; Herrera *et al.* 2012), was remarkably high (70-80%) (Fig 1).

**Figure 1.**
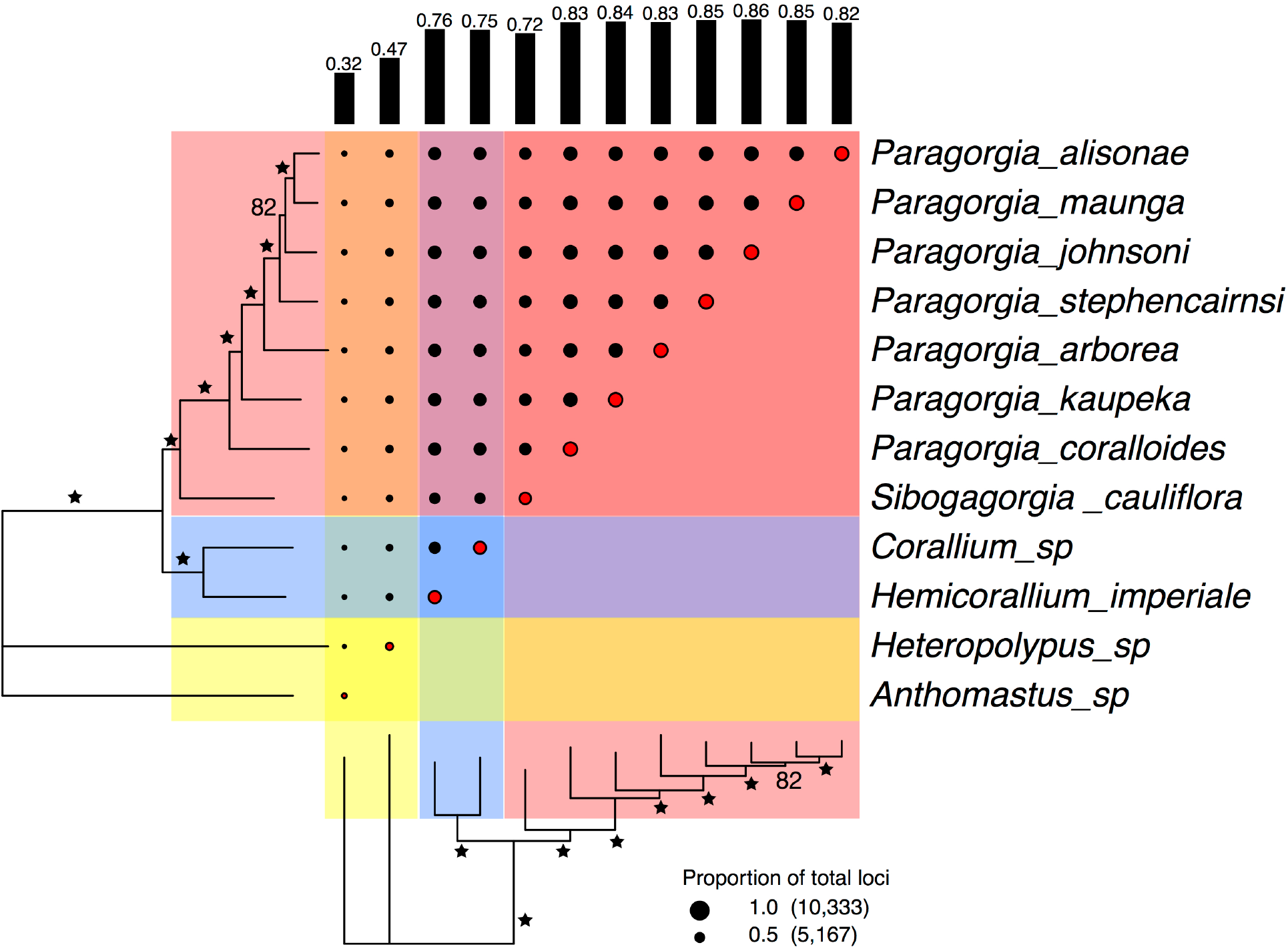
Proportion of loci shared among individuals of the AC clade in the optimal backbone matrix (c 0.80, m 9). Each family is indicated with a different color: red for Paragorgiidae; blue for Coralliidae; and yellow for Alcyoniidae. Black-filled circles represent the proportion of the total number of loci shared among individuals. Red-filled circles represent the proportion of the total number of loci present in each individual. Circle scale shows the number of loci represented by 1.0 and 0.5 circle sizes. Black vertical bars represent the average proportion of loci shared by each individual. Phylogenetic tree was inferred with RAxML. Stars on the tree represent branch bootstrap support of 100. Smaller bootstrap support values are indicated with numbers. This figure was generated with the package RADami (Hipp *et al.* 2014).

We used the selected optimal loci-clustering parameters to generate the **‘PHYLO’** matrix, containing the sequence data of all the 44 octocoral specimens. The use of the parameter value c 0.80 yielded approximately 71 ± 15 thousand loci – with a minimum depth of coverage of 5x and after filtering for paralogs– per specimen (mean depth of clusters used in loci construction was 23 ± 8x) (Table S4). The **PHYLO** matrix contained a total of 5,997 loci that contained data for at least 75% of the specimens (after a second paralog removal). There were 85,293 variable sites in this matrix, of which 53,150 were phylogenetically informative.

### RAD-seq data support a fully resolved phylogeny

The phylogenetic analysis of the **PHYLO** concatenated RAD-seq matrix produced a completely resolved evolutionary tree of the AC specimens (Fig 2). In general, all branches were supported by high (greater than 95) bootstrap values, except for the one supporting the clade of *P. johnsoni, P. alisonae,* and *P. maunga*. Each one of the morphologically identified families, genera, and species in this dataset were monophyletic. The branching pattern of the tree is consistent with an expected transition between coalescent processes among species and genera (long deep branches), and population processes within species (short shallow branches).

**Figure 2.**
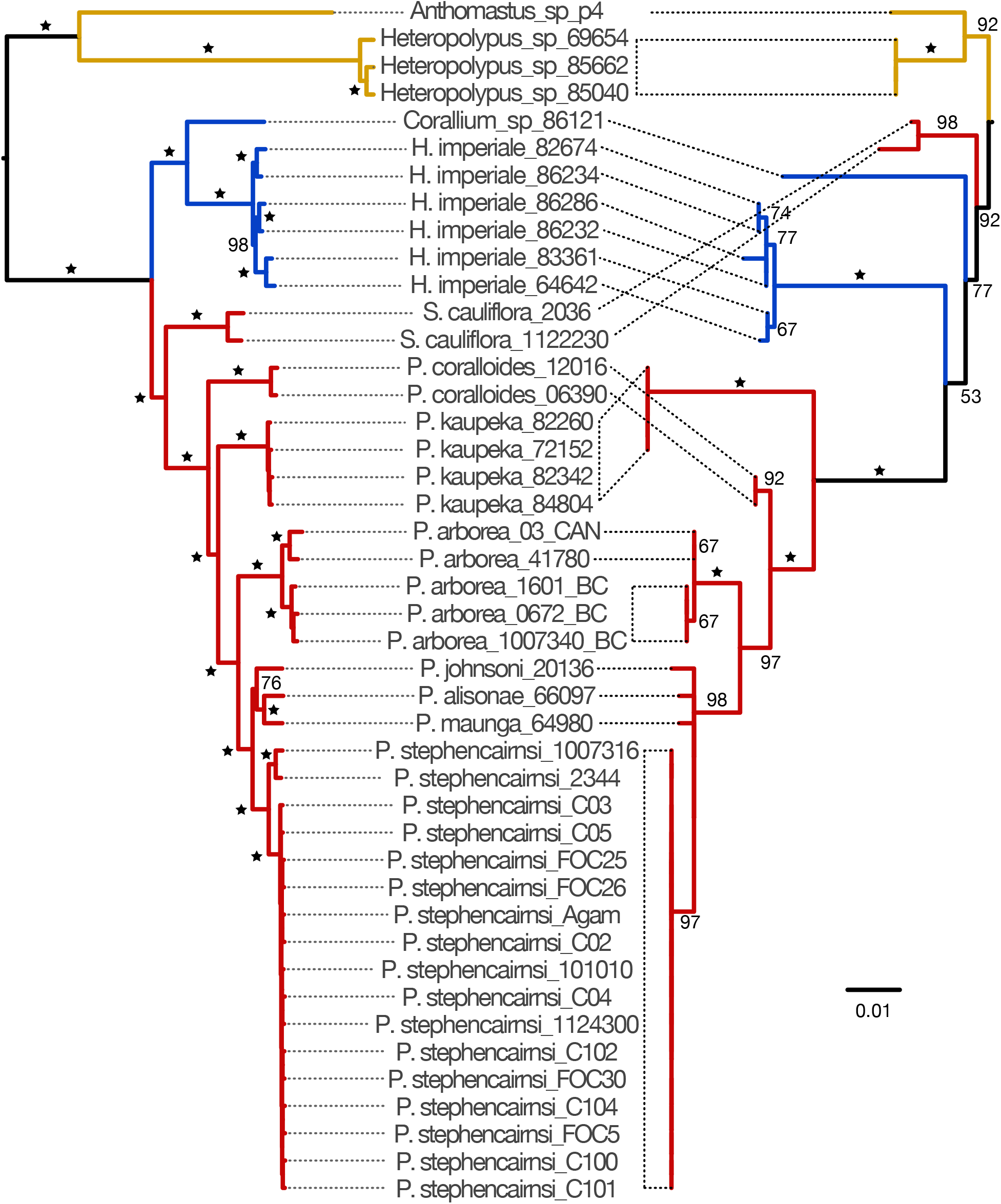
Phylogenetic trees of the AC clade based on RAD-seq and mitochondrial data. Left tree based on the RAD-seq concatenated PHYLO matrix. Right tree based on the mtMutS mitochondrial matrix. Each family is indicated with a different branch color: blue red for Paragorgiidaea; blue for Coralliidae; and yellow for Alcyoniidae. Phylogenetic trees were inferred with RAxML. Stars on the trees represent bootstrap support of 100. Smaller bootstrap values are indicated in numbers. Scale bar indicates substitutions per site.

The topology of the tree obtained with a traditional ‘**mitochondrial’** matrix (711 base pairs of the *mtMutS* gene containing 130 variable sites, of which 101 were phylogenetically informative) was incongruent with the **PHYLO** tree (Fig 2). The **mitochondrial** tree indicated a well-supported alternative divergence order for *P. coralloides* and *P. kaupeka* in the *Paragorgia* clade. In addition, the families Paragorgiidae (bubblegum corals) and Coralliidae (precious corals) were not monophyletic. The bubblegum coral genus *Sibogagorgia* appeared more closely related to the precious corals than to the other bubblegum coral genus *Paragorgia,* and the genera *Corallium* and *Hemicorallium* did not form a clade. However, these alternative relationships were not significantly supported by the bootstrap analysis,. Indeed, a substantial proportion of branches on the **mitochondrial** tree were poorly supported (bootstrap values smaller than 80%).

### RAD-seq data reveal cryptic genetic diversity

Branch-length differences among individuals, as well as well-supported sub-clades, revealed intraspecific genetic diversity that was undetected by the **mitochondrial** matrix. Two sub-clades were revealed by the phylogenetic analysis of the **PHYLO** matrix in the *P. arborea* and *P. stephencairnsi* clades. The sub-clades in *P. arborea* correspond to a pattern of segregation by geographic location with specimens from the north Pacific in one sub-clade, and specimens from the south Pacific and north Atlantic in the other. Contrastingly, the sub-clades in *P. stephencairnsi* correspond to a pattern of segregation by depth with specimens collected shallower than 350m in one sub-clade, and specimens collected deeper than 1000m in the other.

### Current morphological species delimitation is rejected

To evaluate the utility of RAD-seq to perform objective species delimitations in octocorals we focused on specimens the genus *Paragorgia* as it was the best-sampled taxon in our dataset, both in terms of geographic representation and number of morphological species. We used the Bayes Factor Delimitation method with genomic data (BFD*) (Leache *et al.* 2014), which allows for the comparison of conflictive species delimitation models in an explicit multispecies coalescent framework using genome-wide single nucleotide polymorphism (SNP) data. We calculated marginal likelihoods of taxonomy-guided and taxonomy-unguided species delimitation models from a matrix of unlinked SNPs including only specimens of *Paragorgia* (**‘PARAGORGIA’** matrix containing 1,203 SNPs present in all individuals). We compared the marginal likelihood estimates of alternative species delimitation models to the null model ‘**morphid’**, which is based on current morphological species descriptions, using Bayesian factors.

The null model, **morphid,** was rejected in favor of alternative species delimitation models for *Paragorgia* (Fig 3) (**morphid** was ranked 7^th^ among 10 evaluated models in terms of the marginal likelihood estimate). The ‘**PABSTE’** model, which proposes 9 species based on the 7 morphological species in the dataset plus splits corresponding to the sub-clades in *P. arborea* and in *P. stephencairnsi,* received decisive support from Bayes factors as the best species delimitation model. The taxonomy-unguided model ‘**geo’** – which splits the specimens based on the geographic location where they were collected – and the models proposed by the Poisson tree processes (PTP) method based on the **mitochondrial** data matrix, were the lowest ranked and most strongly rejected models overall.

**Figure 3.**
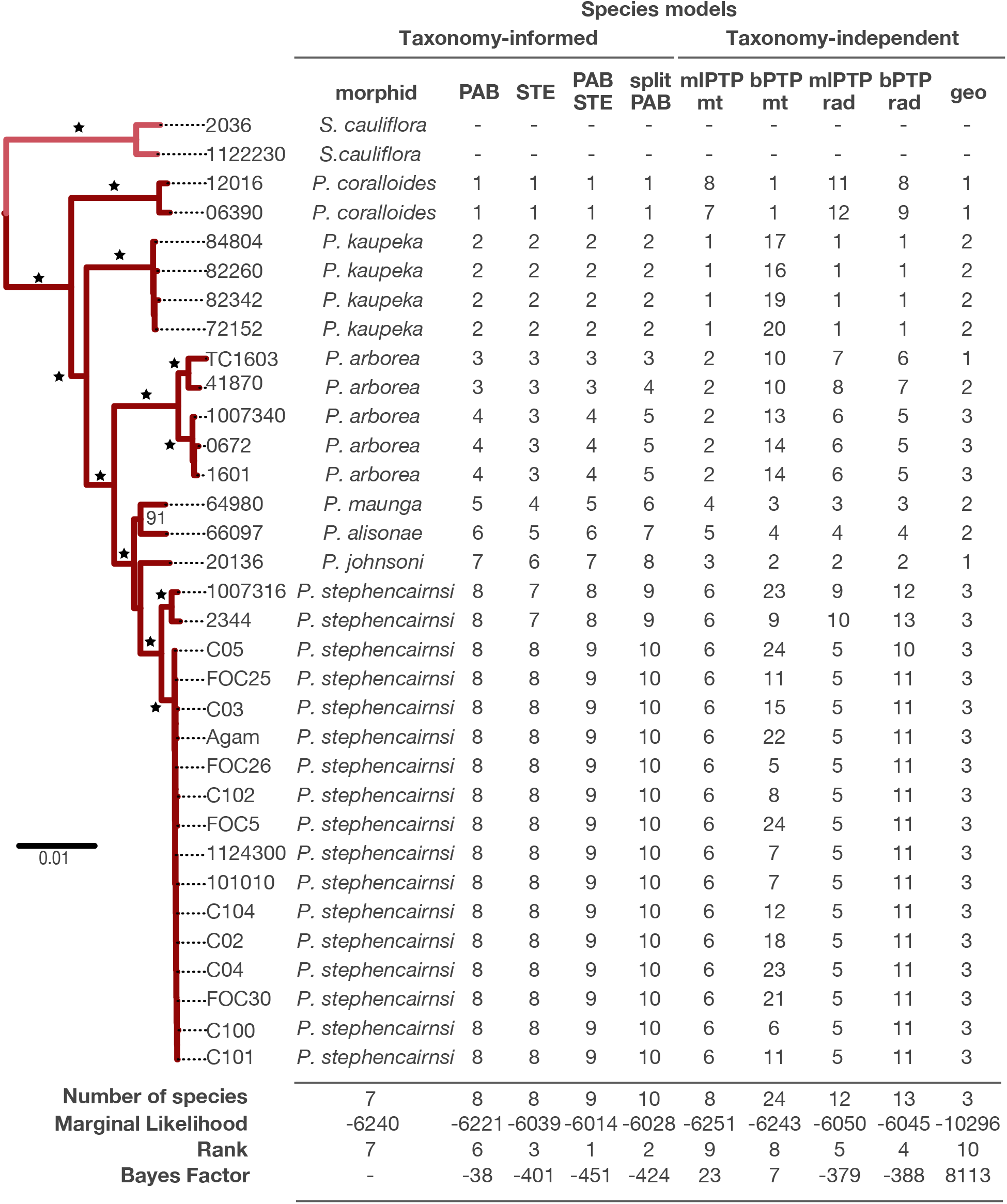
Species delineation hypotheses for *Paragorgia*. Table shows the different species delimitation models for *Paragorgia* evaluated with the BFD* method and their results. *Sibogagorgia* was included as outgroup to root the inferences for *Paragorgia*. Each row indicates a different specimen. Each column indicates a different species delimitation model. The first column, model morphid, indicates the species identifications based on morphology. For all other models, numbers indicate the species assignments. Bottom rows show the total number of species proposed, the marginal likelihood estimate, and rank for each model. The Bayes factor comparisons were calculated with respect to the null morphid model. Phylogenetic tree on the left, shown only for visual reference, was inferred with the RAD-seq concatenated PARAGORGIID matrix in RAxML. Each genus is indicated with a different branch color: pink for *Sibogagorgia*; and dark red for *Paragorgia*. Stars on the trees represent bootstrap support of 100. Smaller bootstrap values are indicated in numbers. Scale bar indicates substitutions per site.

### Concatenated and coalescent species tree analyses are congruent

The topology of the species tree inferred using the SNP **PARAGORGIA** matrix was entirely congruent with the topology generated by the maximum likelihood phylogenetic analysis of the concatenated sequence matrices (Fig 4). The species tree analysis also greatly improved support for the clade of *P. johnsoni, P. alisonae,* and *P. maunga.* The posterior distribution of species trees indicated a small fraction of conflictive topologies concentrating in this region of the tree.

**Figure 4.**
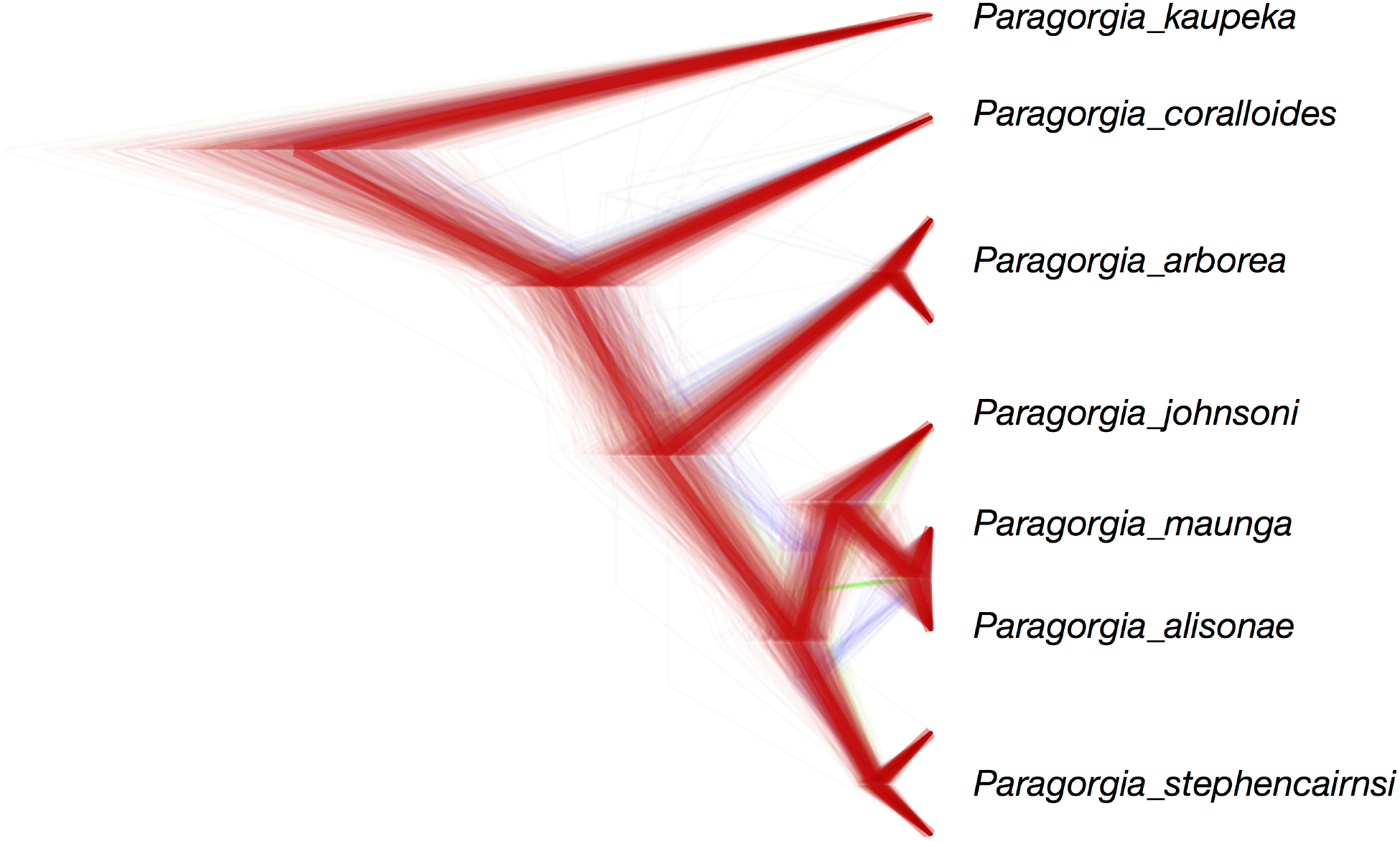
Species tree of *Paragorgia.* This claudogram illustrates the posterior distribution of species trees inferred with SNAPP based on the best species delimitation model PABSTE. High color density is indicative of areas in the species trees with high topology agreement. Different colors represent different topologies. The maximum clade credibility species tree is shown with thicker branches. Trees with the same topology as the maximum clade credibility species tree are colored in red. Trees with different topologies are colored green or blue. With the exception of the branch leading to the clade of *P. johnsoni, P. maunga*, and *P. alisonae,* which has a posterior probability of 0.87, all interior branches have posterior probabilities of 1.0.

## DISCUSSION

### RAD sequencing enables unprecedented phylogenetic resolution

Our analyses of RAD-seq data provide a robust phylogenetic hypothesis for the recalcitrant octocorals in the *Anthomastus-Corallium* clade, a result never achieved before. Moreover, this study, together with the work by Pante *et al.* (2014) in the octocoral genus *Chrysogorgia*, constitute the first applications of RAD-sequencing for phylogenetics and species delimitation in cnidarians. Only a handful of previous studies, using traditional mitochondrial data and the ITS2 and 28S nuclear markers, have attempted to evaluate phylogenetic relationships in the octocoral AC clade (Herrera *et al.* 2010; Ardila *et al.* 2012; Brockman & McFadden 2012; Herrera *et al.* 2012; McFadden & van Ofwegen 2013; Uda *et al.* 2013; Figueroa & Baco 2014). These studies find support for the monophyly of the genus *Paragorgia,* the family Coralliidae, and the sister relationship between the Paragorgiidae and Coralliidae. However, those data do not provide enough phylogenetic resolution to infer the evolutionary relationships among many of the putative morphological species. Furthermore, significant incongruences between mitochondrial and nuclear ITS2 gene trees from AC taxa have been documented (Herrera *et al.* 2010). Here we reproduce similar incongruences when comparing the trees inferred from mitochondrial and RAD-seq datasets (Fig 2). Likewise, Pante *et al.* (2014) documented marked incongruence between trees inferred from mitochondrial and RAD-seq data in *Chyrsogorgia.* These observations suggest that processes that can cause gene tree heterogeneity, such as incomplete lineage sorting and horizontal gene transfer (Maddison 1997; Edwards 2009), may be more prevalent in octocorals than previously recognized.

All of our analyses based on RAD-seq matrices – varying in taxon coverage, degree of divergence among taxa, proportion of missing data, number of loci, and analysis type (concatenated or species tree) – produced completely congruent trees, which together provide extremely high confidence on the phylogenetic hypothesis inferred for the octocoral AC clade (Figs 1, 2 and 3). Consequently, we suggest that single marker gene trees in octocorals, particularly from the mitochondria, should not be considered as robust hypotheses of true species phylogenies on their own, without further validation by multiple informative and independent nuclear loci. We urge systematists to be conservative when making taxonomic rearrangements based on inferences from single-marker data alone.

### RAD-seq data is suitable for phylogenetic inference in divergent taxa

Contrary to the currently accepted idea that RAD-seq data are only suitable for taxa with divergence times younger than 60 million years (MY) (Rubin *et al.* 2012), we demonstrate their suitability well beyond this age threshold. Remarkably, we were able to confidently resolve phylogenetic relationships among genera from different families diverging by at least 80 MY in the AC clade. The split between the families Paragorgiidae and Coralliidae has been dated, using coralliid fossils, to be between 80-150 MY old (Ardila *et al.* 2012; Herrera *et al.* 2012). Park *et al.* (2012) estimated the age of the most recent common ancestor of the Coralliidae at approximately 50 MY (25-100 MY 95% confidence region), using independent cnidarian fossils for molecular clock calibration. The split with the genera *Anthomastus* and *Heteropolypus* is likely older than 100 MY. It is without question that, due to evolution at restriction sites, the number of RAD loci among taxa for which orthology can be established decreases rapidly as divergence increases. However, we suggest that the suitability limits of RAD-seq for phylogenetics in divergent taxa cannot be assessed in terms absolute time, but depend on taxon-specific factors such as mutation rate, generation time and effective population size.

Bioinformatic studies addressing the issue of extent of the suitability of RAD-seq for phylogenetic inference have focused mainly on *Drosophila* as study model (Rubin *et al.* 2012; Cariou *et al.* 2013). Longer generation times and lower metabolic rates in taxa like deep-sea corals, relative to those in organisms like *Drosophila,* could cause a reduction in mutation rates (see review by Baer *et al.* (2007)), which may in turn decrease the evolutionary rates at restriction sites and allow for phylogenetic inferences using RAD-seq in situations of deeper divergence. Consistent with this hypothesis, we observe a nucleotide diversity (π) calculated across all octocoral specimens from the **PHYLO** matrix of 0.012 ± 0.002 (considered a minimum since RAD-seq can underestimate diversity (Arnold *et al.* 2013); see Table S5 and Table S6 for individual values), which is significantly lower than the nucleotide diversity in many of the *Drosophila* species included in the bioinformatic studies by Cariou *et al.* (2013) and Rubin *et al.* (2012). Nonetheless, there are other important factors known to influence genetic diversity across species – and likely the evolutionary rate as well. These factors include the effective population size, selection, habitat kind, geographic range, and mating system (Leffler *et al.* 2012). To sum up, we argue that RAD-seq can be successfully used to infer phylogenetic relationships in certain taxa with deeper divergences than previously suggested. This is particularly true when the number of RAD loci is maximized through the choice of restriction enzymes with higher cutting frequencies in the target taxon (Herrera *et al.* 2014).

### RAD-seq allows the formulation of robust species delineations

Our study, the first statistical rigorous test of species hypothesis in octocorals, provides conclusive evidence rejecting the current morphological species delimitation model for the genus *Paragorgia*. We find decisive support for a nested model that combines species boundaries from morphological taxonomy with cryptic diversity linked to environmental variables of geographic location and depth (Figs 3 and 4). This nested model, proposes 9 species among the examined specimens. Five of these species correspond to the morphological species *P. coralloides, P. kaupeka, P. alisonae, P. johnsoni,* and *P. maunga.* Two splits, corresponding to sub-clades in the morphological species *P. arborea* and in *P. stephencairnsi*, indicate cases of cryptic species.

Herrera *et al.* (2012) found significant genetic differentiation of the north Pacific populations of *P. arborea* relative to the south Pacific, Atlantic and Indian ocean populations, and suggested that these populations may represent sub-species. The north Pacific populations of *P. arborea* were previously defined as a separate species, *P. pacifica,* by Verrill (1922) based on gross colony morphology, but later combined into a single species by Grasshoff (1979). Sánchez (2005) suggested potential small differences in medullar sclerite sizes and ornamentation between north Pacific specimens and specimens form elsewhere. However, we were unable to recognize these morphological differences in the few examined specimens in this study. Nonetheless, based on the decisive support for the split of *P. arborea* from analysis of genome-wide SNP makers indicates, we resurrect the species *Paragorgia pacifica* for the north Pacific populations of formerly *P. arborea*. We find no evidence of cryptic speciation between the north Atlantic and south Pacific *P. arborea* and therefore conclude it should be considered a single species as previously suggested by Herrera *et al.* (2012).

Depth is an important factor contributing to genetic differentiation and formation of species in the ocean, both shallow (Carlon & Budd 2002; Prada & Hellberg 2013) and deep (Miller *et al.* 2011; Jennings *et al.* 2013; Quattrini *et al.* 2013; Glazier & Etter 2014). The observed cryptic differentiation between specimens of *P. stephencairnsi* collected shallower than 350m and deeper than 1000m indicates that depth is also a diversifying force in octocorals from the AC clade, which had gone undetected due to the low variability of traditional sequence data (Herrera *et al.* 2012). The holotype of *P. stephencairnsi* was collected from approximately 350m in the Georgia Strait of British Columbia, overlapping in depth range and geographic region with that of most of the specimens from the shallow sub-clade examined in this study. Therefore, we propose to conserve that name *P. stephencairnsi* for that shallow sub-clade, and consider the deep sub-clade as a new species.

Other recent species delimitation studies in anthozoan corals have also revealed significant incongruences when comparing morphological and single-locus species delimitation hypotheses (particularly from mitochondrial data) with phylogenetic evidence from multi-locus datasets (Pante *et al.* 2014; Prada *et al.* 2014). In line with the findings of Pante *et al.* (2014), we find that specimens of *Paragorgia* sharing identical *mtMutS* haplotypes can belong to more than one species. Contrastingly, Herrera *et al.* (2012) present strong evidence showing significant mitochondrial haplotype diversity in the south Pacific and north Atlantic populations of *Paragorgia arborea*. Our observations, together with those from the aforementioned studies, constitute compelling evidence indicating that there is no solid basis for the widespread assumption that *mtMutS* haplotypes may be equivalent to individual octocoral species, as proposed by Thoma *et al.* (2009). The analysis with RAD-seq, or alternative genomic multi-locus methods, of a larger number of specimens from diverse geographic locations and depth horizons will likely reveal further cryptic diversity not characterized by mitochondrial haploytypes (see Fig S2, Fig S3, and Table S8), and thus further illuminates taxonomy and systematics in this an other groups.

## CONCLUSIONS

In this case study we demonstrate the empirical utility of RAD-seq to resolve phylogenetic relationships among divergent and recalcitrant taxa and to objectively infer species boundaries by testing alternative delimitation hypotheses. We were able to make use of RAD-seq to overcome long-standing technical difficulties in octocoral genetics, and to resolve fundamental questions in species definitions and systematics. We show that classic morphological taxonomy can greatly benefit from integrative approaches that provide objective tests to species delimitation hypothesis. Our results pave the way for addressing further questions in biogeography, species ranges, community ecology, population dynamics and evolution of octocorals and other marine taxa. The results from this study also represent a valuable reference resource for the development of tools, such as SNP arrays, that can be used to perform accurate species identifications, and generate species inventories that will aid the design and implementation of conservation and management policies.

## METHODS

To perform identifications using current morphological species descriptions we performed colony observations and scanning electron microscopy of sclerites on 44 octocoral specimens from the AC clade (Table S1).

To obtain a genome-wide set of markers that could be useful for phylogenetic inferences of deep-divergent taxa and species delimitation in the AC clade (greater than 100 million years) we performed RAD sequencing with a the 6-cutter restriction enzyme PstI, which is predicted to cut between 32,000 and 110,000 times in the genome of an octocoral (Table S7). This predicted range was obtained using the observed frequency of the PstI recognition sequence, and its probability calculated using a trinucleotide composition model, in the genomes of the cnidarians *Nematostella vectensis, Acropora digitifera, Hydra vulgaris,* and *Alatina moseri* (Herrera *et al.* 2014). Genome size range of 0.3-0.5 pg was used based on observations obtained through flow cytometry in gorgoniid octocorals by Luisa Dueñas at the Universidad de los Andes, Bogotá, Colombia (personal communication). Total genomic DNA was purified from specimens following protocols described in Herrera *et al.* (2015). Concentration-normalized genomic DNA was submitted to Floragenex Inc (Eugene, OR). for library preparation and RAD sequencing. Libraries were sequenced by 48-multiplex, using 10-base pair barcodes, on a single lane of an Illumina Hi-Seq 2000 sequencer.

To compare the inferences obtained from RAD-seq data with the inferences drawn from traditional genetic barcoding data, we performed targeted sequencing of the mitochondrial *mtMutS* gene — a genetic marker widely used for phylogenetics and species delimitation studies in octocorals. Polymerase chain reactions were carried out following the protocols by Herrera *et al.* (2015). Primer pairs used for amplifications were AnthoCorMSH (Herrera *et al.* 2010) and Mut-3458R (Sánchez *et al.* 2003). Negative controls were included in every experiment to test for contamination. Purified PCR products were submitted to Eurofins Genomics (Eurofins MWG Operon, Inc.) for sequencing.

### RAD-seq data filtering

Sequence reads were de-multiplexed and quality filtered with the process_radtags program from the package Stacks v1.20 (Catchen *et al.* 2013). Barcodes and Illumina adapters were excluded from each read and length was truncated to 91bp (-t 91) Reads with ambiguous bases were discarded (-c). Reads with an average quality score below 10 (-s 10) within a sliding window of 15% of the read length (-w 0.15) were discarded (-r). The rescue barcodes and RAD-tags algorithm was enabled (-r). Additional filtering, and the clustering within and between individuals to identify loci was performed using the program pyRAD v2.01 (Eaton 2014). Reads with more than 33 bases with a quality score below 20 were discarded.

### RAD-seq loci clustering and phylogenetic inference

We investigated different combinations of clustering thresholds (c 0.80, 0.85 and 0.90) and minimum number of taxa per locus (m 4, 6, and 9) in a reduced dataset that included one individual from each of the 12 putative morphological species. The minimum depth of coverage required to build a cluster and the maximum number of shared polymorphic sites in a locus were kept constant at 4 (d) and 3 (p) respectively. Loci sequences were concatenated into combined matrices. We refer to these 9 resulting matrices as the **‘backbone’** matrices. Each of the resulting backbone matrices was analyzed in RAxML-HPC2 v8.0 (Stamatakis 2006) for maximum likelihood (ML) phylogenetic tree inference. For this, and all the other phylogenetic analyses in RAxML, we assumed a generalized time-reversible DNA substitution model with a gamma-distributed rate variation across sites (GTR GAMMA). Branch support was assessed by 500 bootstrap replicates.

We selected an optimal combination of loci clustering parameters as the set of parameters that minimized the number of missing data and maximized the number of phylogenetically informative sites while producing a highly supported phylogenetic tree. The optimal set of parameters chosen was a clustering threshold of 80% similarity among sequences (c 0.80) and a minimum coverage of taxa per locus of 75% (m 9). A concatenated matrix containing the sequence data of all the 44 octocoral specimens, denominated **‘PHYLO’,** was built using this parameter combination (c 0.80, m 33) in pyRAD and subsequently analyzed in RAxML.

### Phylogenetic inference with traditional genetic barcoding data

To compare the tree topology obtained from the phylogenetic inferences of the **PHYLO** RAD-seq dataset with traditional genetic barcoding data we analyzed the ‘**mitochondrial’** dataset (containing the *mtMutS* sequences) using RAxML. These two datasets – **PHYLO** and **mitochondrial** – contain data from the same individuals. To place the specimens from this study in a broader phylogenetic context we also analyzed the **mitochondrial** dataset in RAxML with the addition of *mtMutS* data from 233 additional specimens belonging to the AC clade, as well as outgroups (see Table S8, Fig S2, and Fig S3).

### Testing species delimitation models for Paragorgia

We constructed 5 taxonomy-guided species delimitation models for *Paragorgia*: **i**) ‘**morphid’** model: 7 species based on current morphological species descriptions (Sánchez 2005); **ii**) ‘**PAB’** model: 8 species based on the 7 morphological species plus a split of *P. arborea* based on previous evidence of genetic differentiation of north Pacific populations (Herrera *et al.* 2012); **iii**) ‘**STE’** model: 8 species based on the 7 morphological species plus a split of *P. stephencairnsi* based on depth differences (specimens collected <350m vs. >1000m), as depth is known to be an important structuring variable in marine taxa (Jennings *et al.* 2013; Prada & Hellberg 2013; Quattrini *et al.* 2013); i**v**) ‘**PABSTE’** model: 9 species based on the 7 morphological species plus the splits of the **PAB** and **STE** models; **v**) ‘**splitPAB’** model: 10 species based on the 7 morphological species plus the split of the **STE** model and an additional split in the **PAB** model where *P. arborea* is split in 3 species corresponding to the ocean basin where the specimens were collected (north Pacific, south Pacific and north Atlantic).

We also generated taxonomy-unguided species delimitation models for *Paragorgia* through Bayesian and ML implementations of the Poisson tree processes model (PTP) (available at http://species.h-its.org/ptp/). PTP estimates the number of speciation events in a rooted phylogenetic tree in terms of nucleotide substitutions (Zhang *et al.* 2013). We used PTP to analyze the trees obtained from phylogenetic inferences in RAxML of reduced *mtMutS* and RAD-seq datasets that include only members of the family Paragorgiidae (genera *Paragorgia* and *Sibogagorgia*). The ‘**PARAGORGIIDAE’** RAD-seq concatenated matrix was generated in pyRAD using a clustering threshold of 80% similarity among sequences (c 0.80) and a minimum coverage of taxa per locus of 100% (m 33). The resulting phylogenetic trees of *Paragorgia* were rooted with the specimens of *Sibogagorgia* and analyzed by the PTP method using a Markov Chain Monte Carlo (MCMC) chain length of 500,000 generations (100 thinning, 25% burnin). We assessed convergence by examining the likelihood trace. The combinations of the ML or Bayesian PTP implementations (mlPTP and bPTP, respectively) with the *mtMutS* or RAD-seq trees of *Paragorgia* resulted in four species delimitation models: **i) ‘mlPTP*mt’*** model; **ii) ‘bPTP*mt’*** model; **iii) ‘mlPTPrad’** model; and **iv) ‘bPTPrad’** model. Lastly, because deep-sea corals are known to show genetic differentiation at ocean basin/regional scales (Miller *et al.* 2011; Morrison *et al.* 2011; Herrera *et al.* 2012), we constructed an additional taxonomy-unguided species delimitation model – the ‘**geo’** model – based on the geographic location where the specimens were collected (north Pacific, south Pacific or north Atlantic ocean basins).

To estimate the marginal likelihood of each species delimitation model we generated a matrix including only specimens of *Paragorgia*, denominated **‘PARAGORGIA’** using a clustering threshold of 80% similarity among sequences (c 0.80) and a minimum coverage of taxa per locus of 100% (m 31) in pyRAD. In contrast to the **backbone**, **PHYLO**, and **PARAGORGIIDAE** RAD-seq matrices, this matrix contained the data of one SNP per locus and not the entire locus sequence. We analyzed these data using the implementation of BFD* in the SNAPP (Bryant *et al.* 2012) plug-in for the program BEAST v2.1.3 (Bouckaert *et al.* 2014). We performed a path-sampling of 48 steps, with a MCMC chain length of 100,000 (10,000 pre-burnin), following the guidelines from Leache *et al.* (2014). Bayesian factors were calculated from the marginal likelihood estimates for each model and compared using the framework proposed by Kass and Raftery (1995)

### Species tree inference

To test the tree topology in the genus *Paragorgia* obtained by the phylogenetic analysis of the **PHYLO** and **PARAGORGIIDAE** concatenated matrices we performed a species tree inference from the SNP data in the **PARAGORGIA** matrix using the program SNAPP. This program allows the inference of species trees from unlinked SNP data (only one SNP per locus retained) bypassing the inference of individual gene trees (Bryant *et al.* 2012). We performed 3 independent runs using a MCMC chain length of 10,000,000 (sampling every 1,000 generations; pre-burnin of 1,000) with default prior distributions for coalescence rate, mutation rate and ancestral population size parameters. We assessed convergence to stationary distributions and effective sample sizes >200 after 10% burnin in the program TRACER (Rambaut & Drummond 2007). Species trees in the posterior distribution were summarized with the program DENSITREE v2.01 (Bouckaert 2010).

## ACKNOWLEDGEMENTS

This research was supported by the National Geographic Society/Waitt Foundation (W285-13 to SH); the National Oceanic and Atmospheric Administration (NA09OAR4320129 to TS); the National Science Foundation (OCE-1131620 to TS); the National Aeronautics and Space Administration (NNX09AB76G to TS); and the Academic Programs Office (Ocean Ventures Fund to SH), the Ocean Exploration Institute (Fellowship support to TMS) and the Ocean Life Institute of the Woods Hole Oceanographic Institution (WHOI).

Specimens provided by the National Institute of Water and Atmospheric Research (NIWA) were collected under research programs: Kermadec Arc Minerals, funded by the New Zealand Ministry of Business, Innovation & Employment (MBIE), Auckland University, Institute of Geological and Nuclear Science (GNS), and WHOI; Ocean Survey 20/20 funded by Land Information New Zealand; Impact of resource use on vulnerable deep-sea communities (CO1X0906), funded by MBIE; Nascent Inter-Ridge Volcanic And Neotectonic Activity, funded by the Ministry for Primary Industries (MPI), GNS, MBIE, and the U. of New Hampshire; Scientific Observer Program funded by MPI; and the Joint New Zealand-USA 2005 NOAA Ring of Fire Expedition, part of NIWA’s Seamount Program (FRST CO1X0508). For enabling access to key specimens we thank K. Schnabel (NIWA), S. Mills (NIWA), D. Tracey (NIWA), M. Clark (NIWA), A. Rowden (NIWA), S. Cairns (Smithsonian), E. Cordes (Temple U.), A. Quattrini (Temple U.), G. Workman (Department of Fisheries and Oceans Canada - DFO), M. Wyeth (DFO), K. Anderson (DFO), M. Frey (Royal British Columbia Museum - RBCM), H. Gartner (RBCM), L. Watling (U. Hawaii), J. Adkins (CalTech). We thank P. Aldersdale (CSIRO), N. Ardila (ECOMAR) and J. Sanchez (U. Andes) for assistance with morphological identifications. We also thank E. O’Brien (WHOI), D. Forsman (WHOI), J. Fellows (WHOI), J. & S. Schooner, K. Heylar, and N. McDaniel for invaluable assistance during scuba diving fieldwork in British Columbia (DFO scientific license FIN130270). We thank the chief scientists, masters, crew, scientific personnel, and funding agencies of expeditions AT07-35, KOK0506, Lophelia II 2009, RB-0503, TAN1007, TAN1104, TAN1206, and TAN1213. We also thank A. Tarrant and A. Reitzel for providing helpful comments that improved this manuscript.

## SUPPLEMENTARY MATERIALS

**Figure S1.**
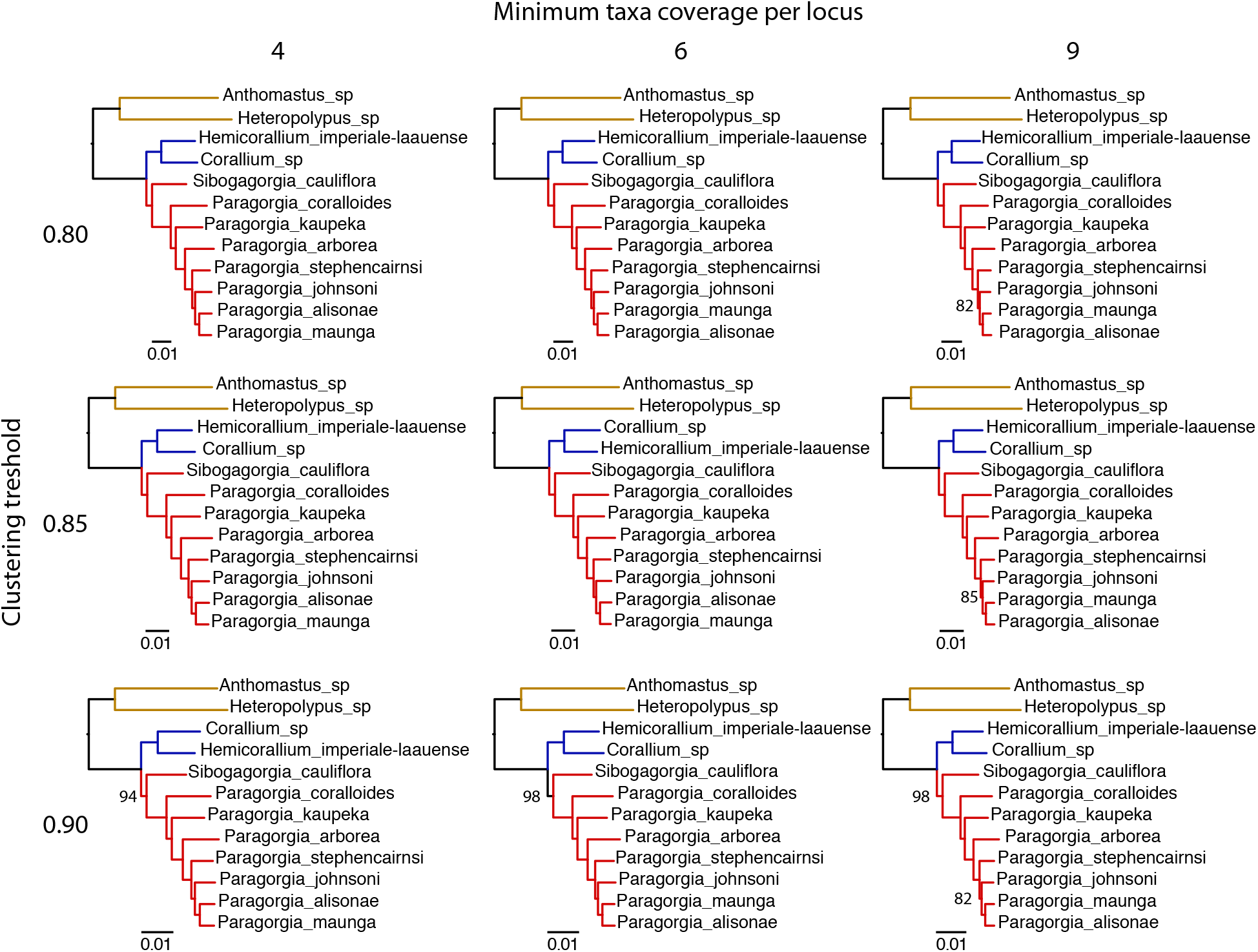
Phylogenetic trees based on backbone matrices. Trees inferred from the 9 backbone RAD-seq matrices built with different parameters of clustering threshold (c 0.80, 0.85 and 0.90; indicated by vertical labels) and minimum number of taxa per locus (m 4, 6, and 9; indicated by horizontal labels). Each family is indicated with a different branch color: red for Paragorgiidae; blue for Coralliidae; and yellow for Alcyoniidae. Trees were inferred with RAxML. All interior branches have bootstrap support values of 100, except for those shown. Scale bars indicate substitutions per site.

**Figure S2.**
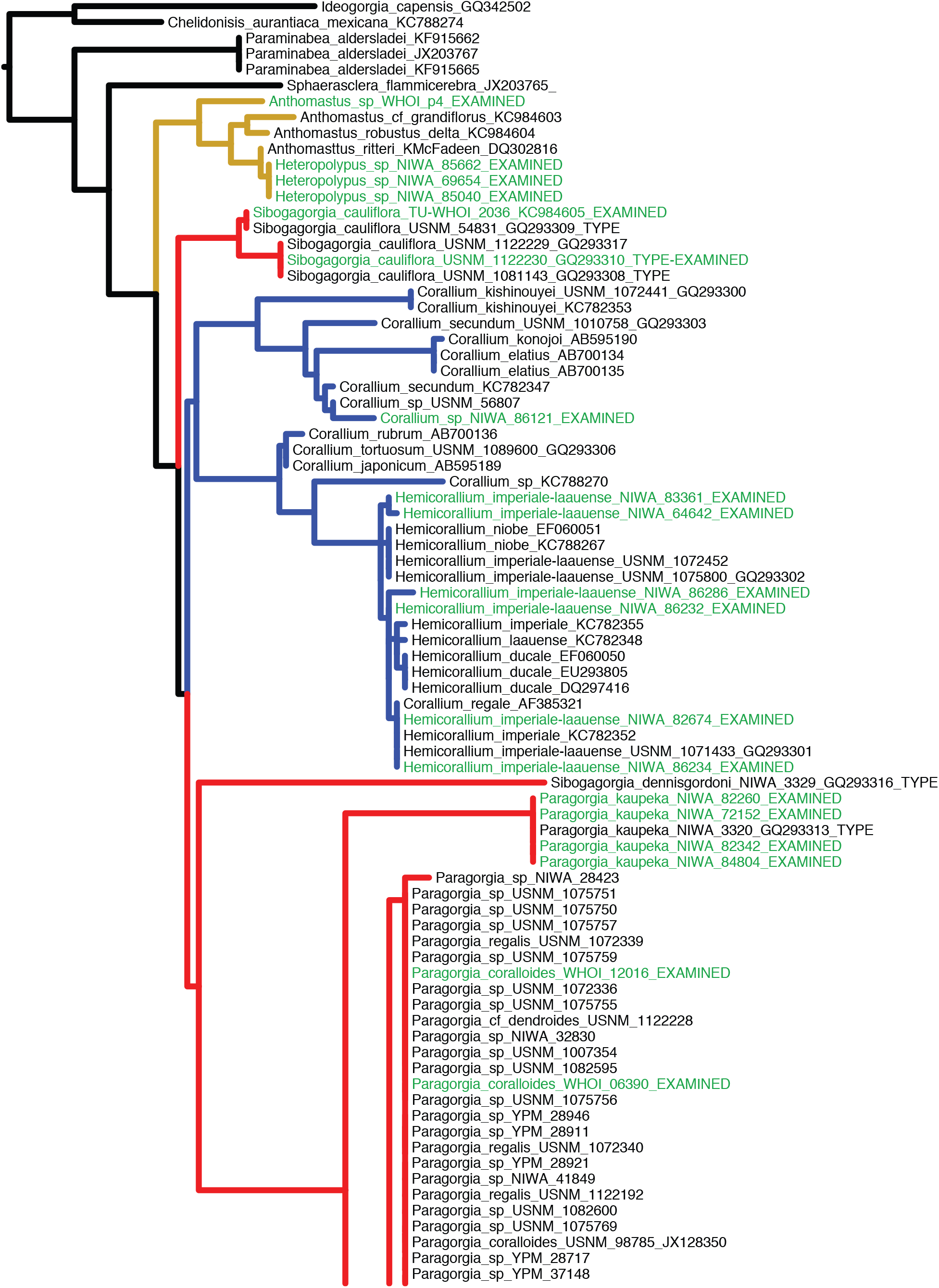
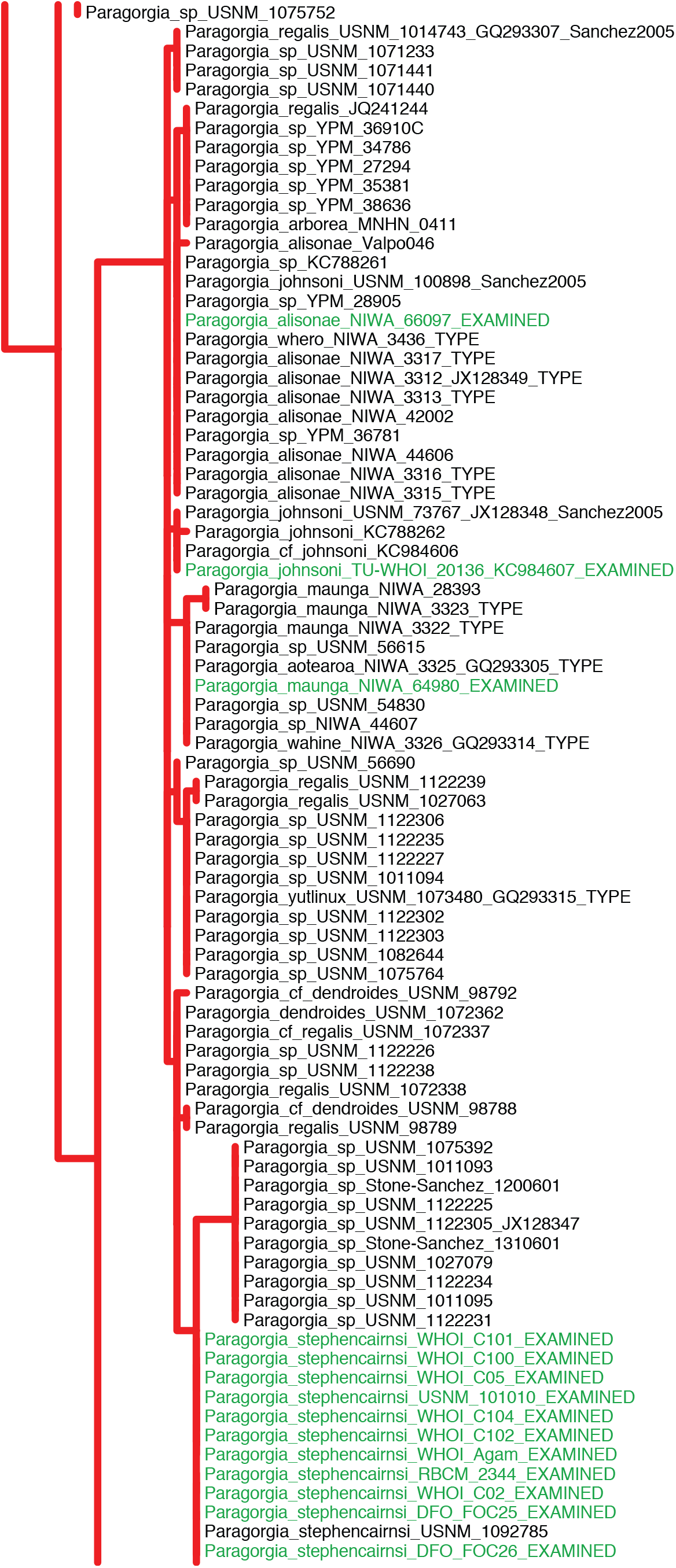
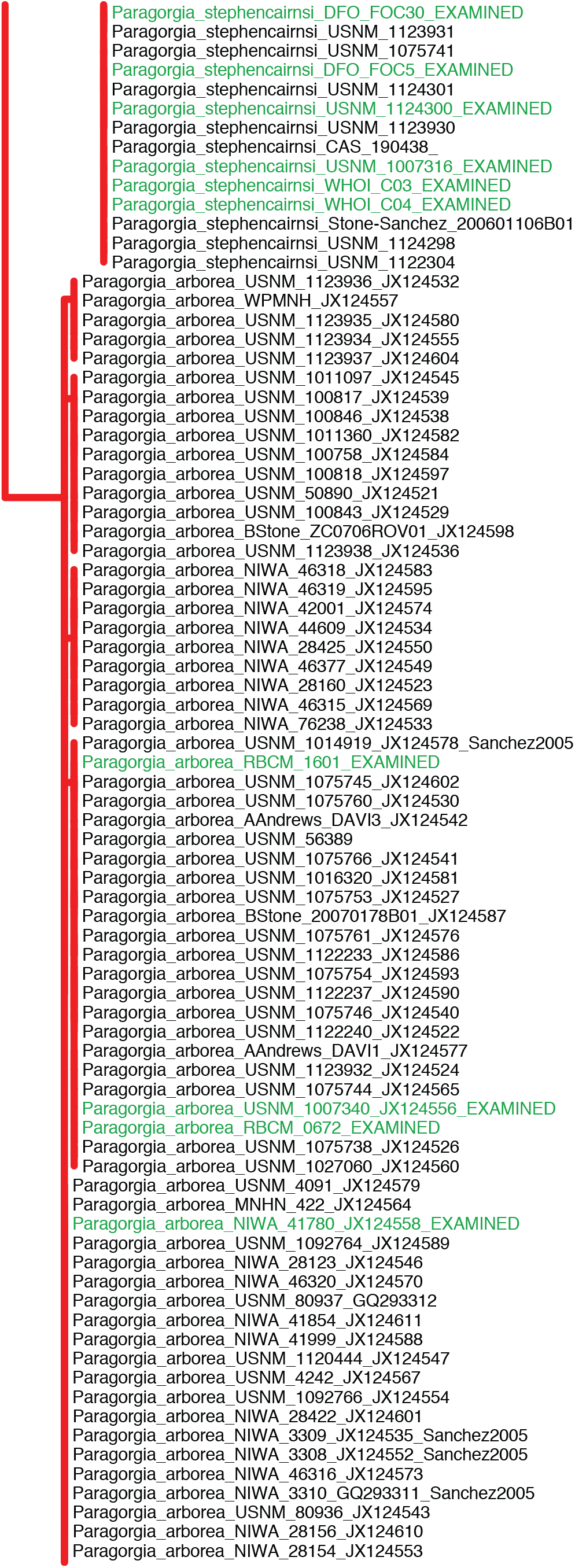
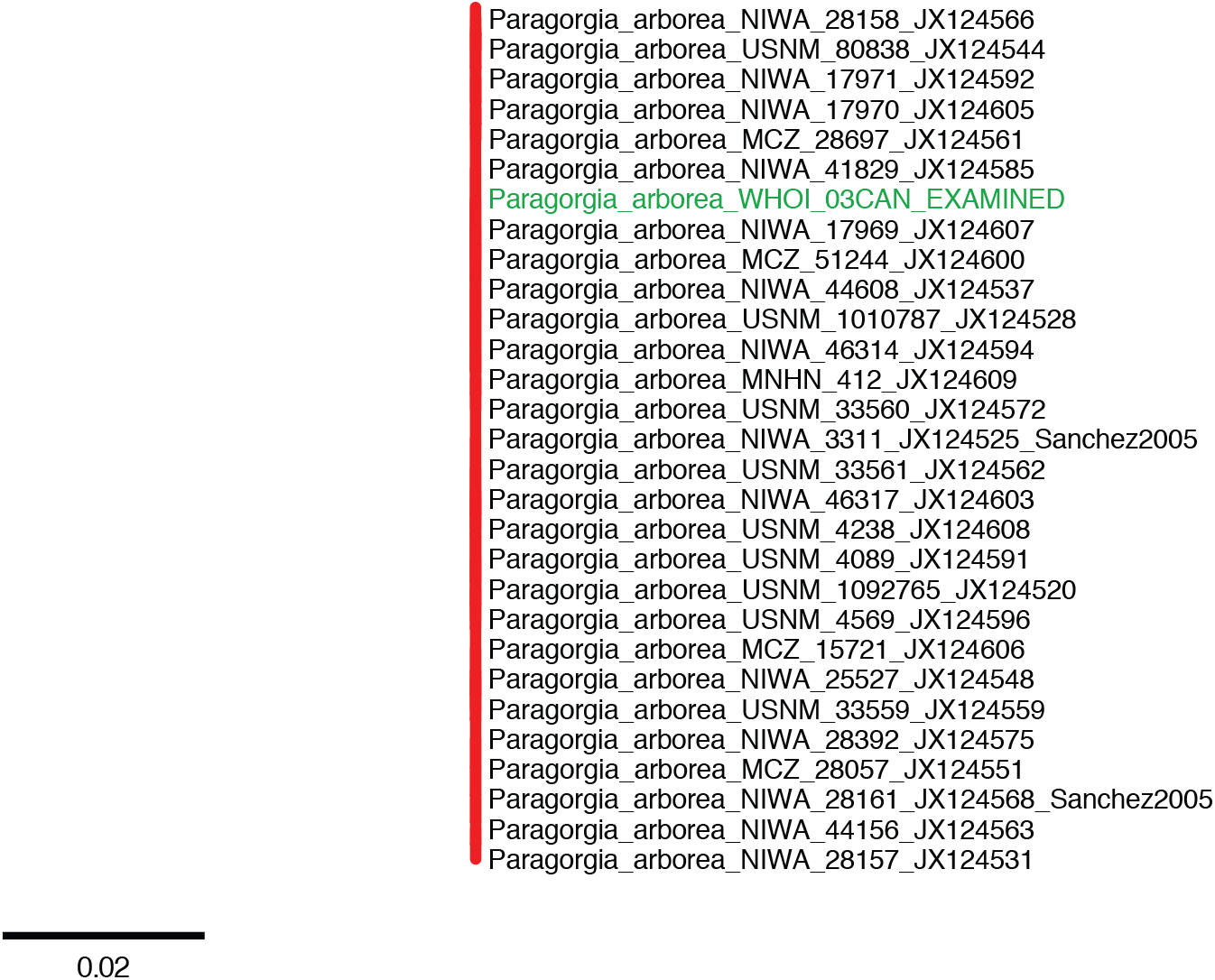
Mitochondrial *mtMutS* gene tree of all available sequences for the clade AC. Tree inferred from *mtMutS* sequence data from specimens examined in this study, GenBank, and additional specimens. Each family is indicated with a different branch color: red for Paragorgiidae; blue for Coralliidae; and yellow for Alcyoniidae. Outgroups are indicated with black branches. Specimens examined in detail in this study are indicated with green labels. Type specimens are labeled TYPE. Specimens examined in Sanchez (2005) are labeled “Sanchez2005”. Tree was inferred with RAxML. Scale bars indicate substitutions per site.

**Figure S3.**
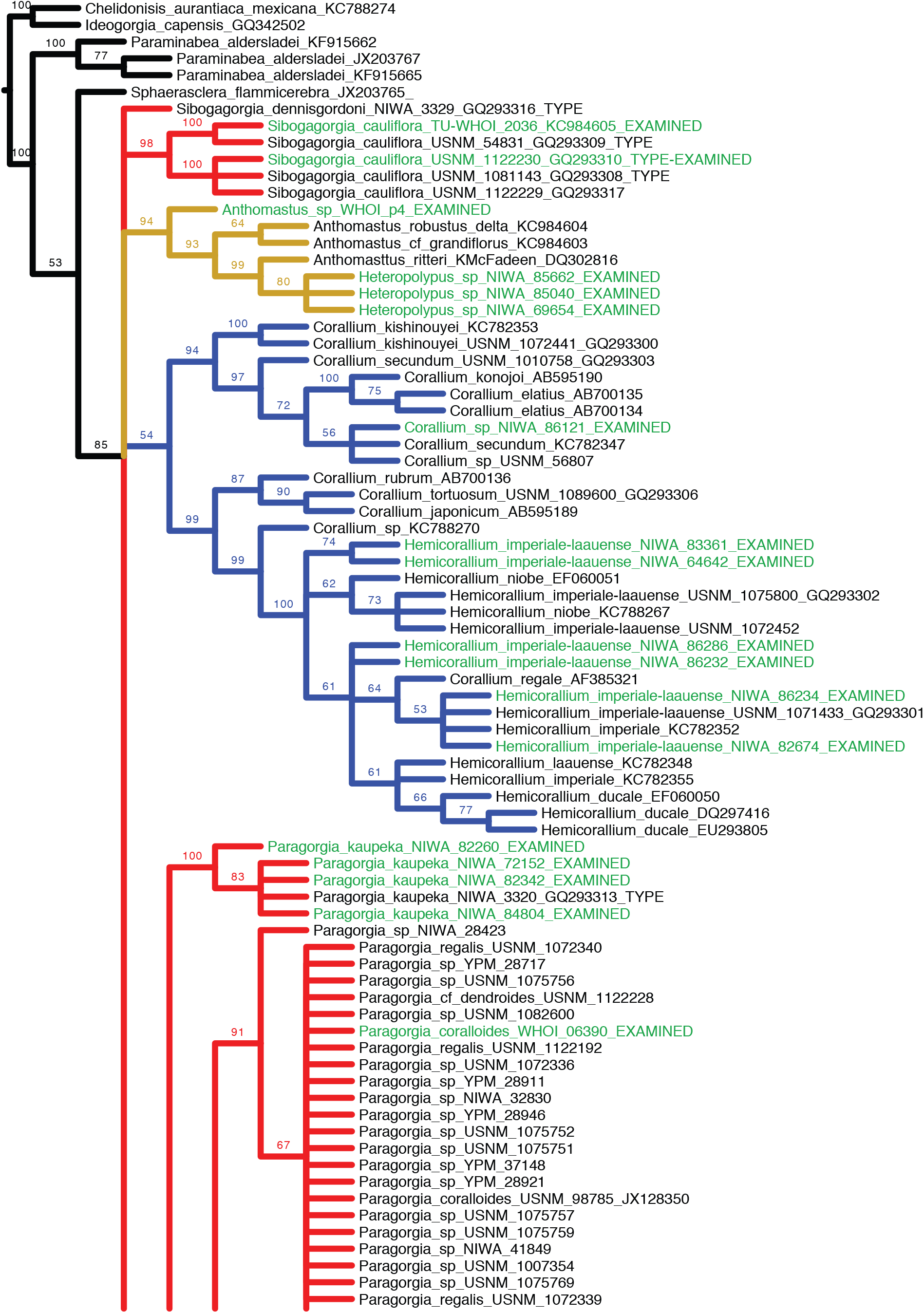
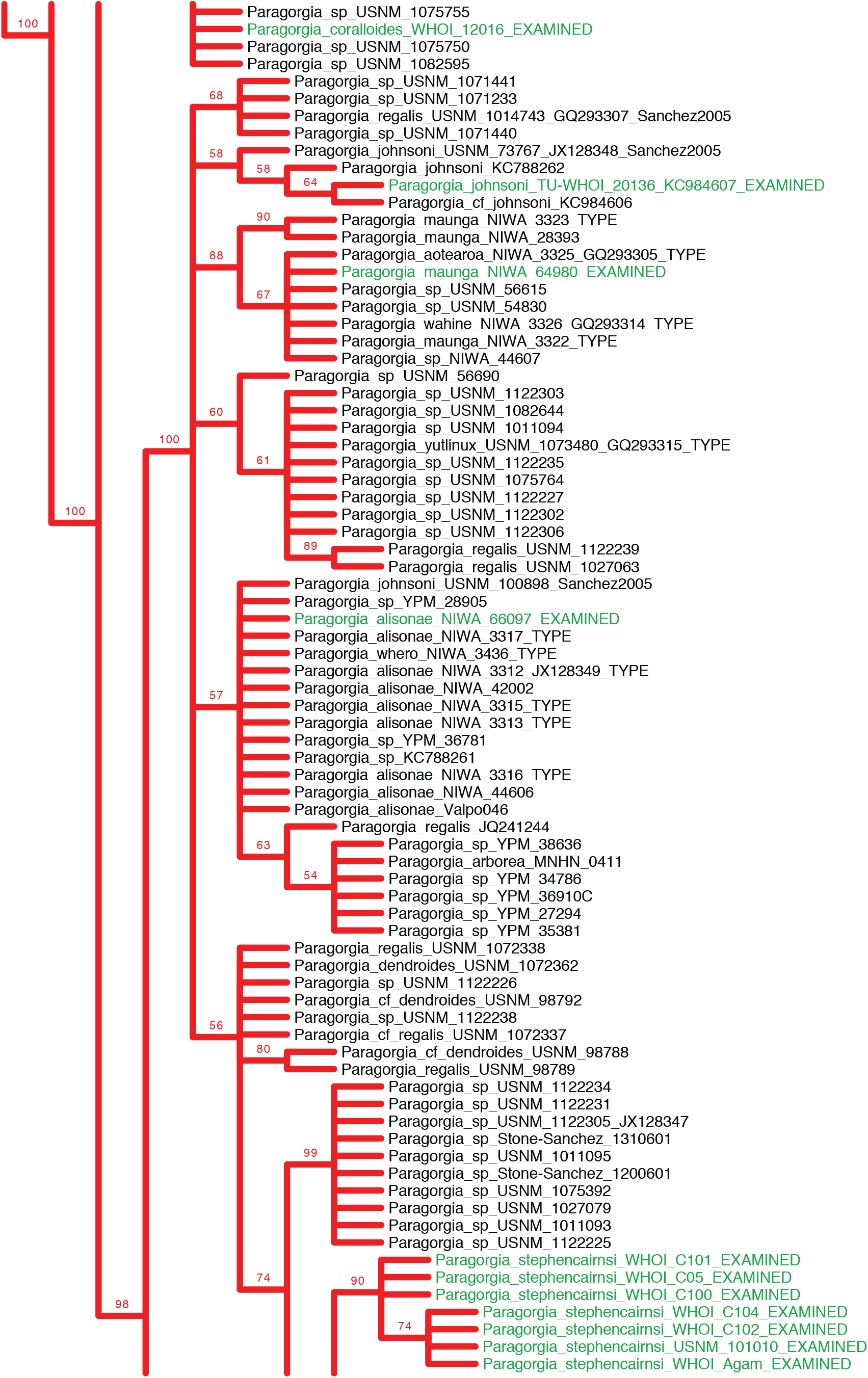
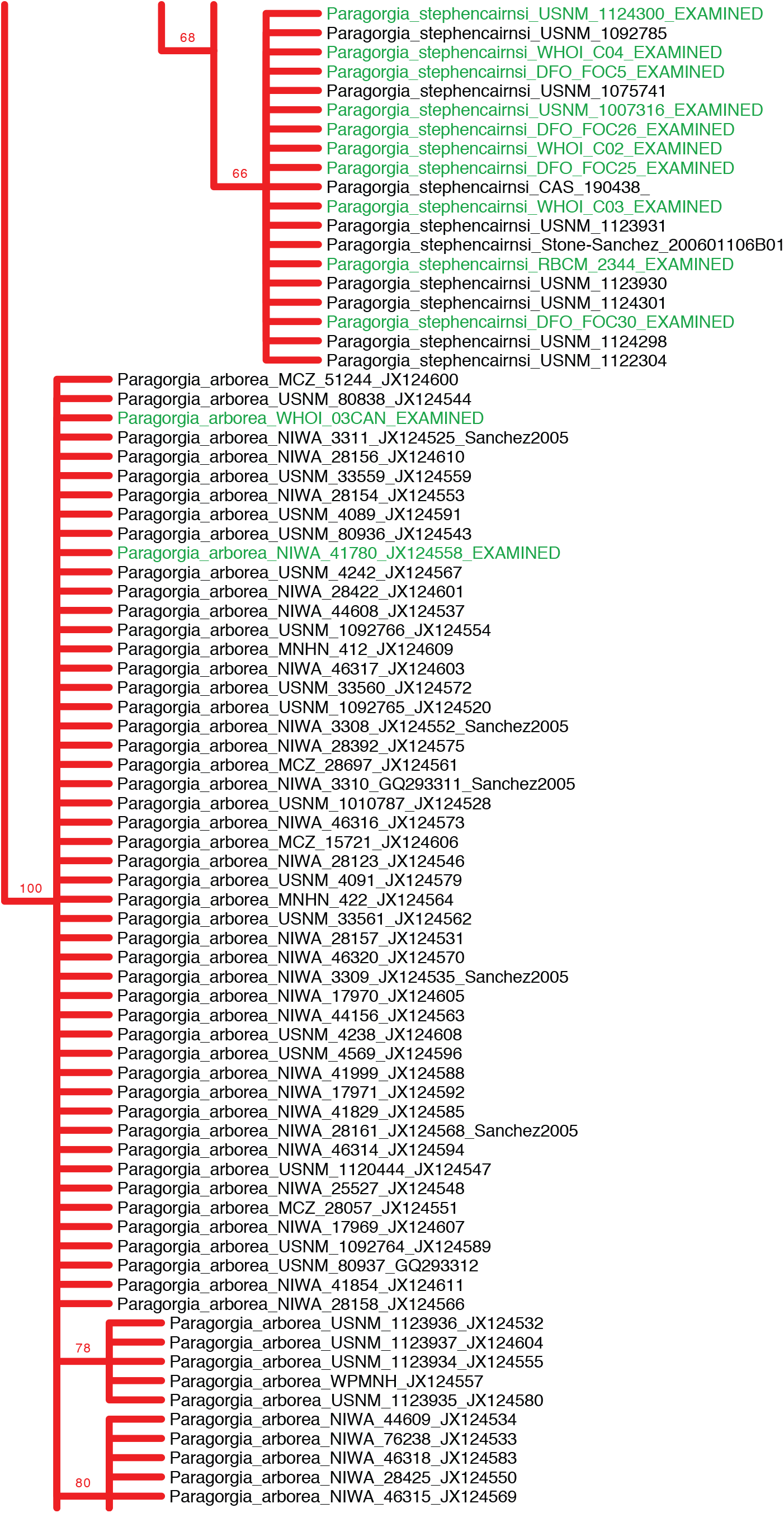
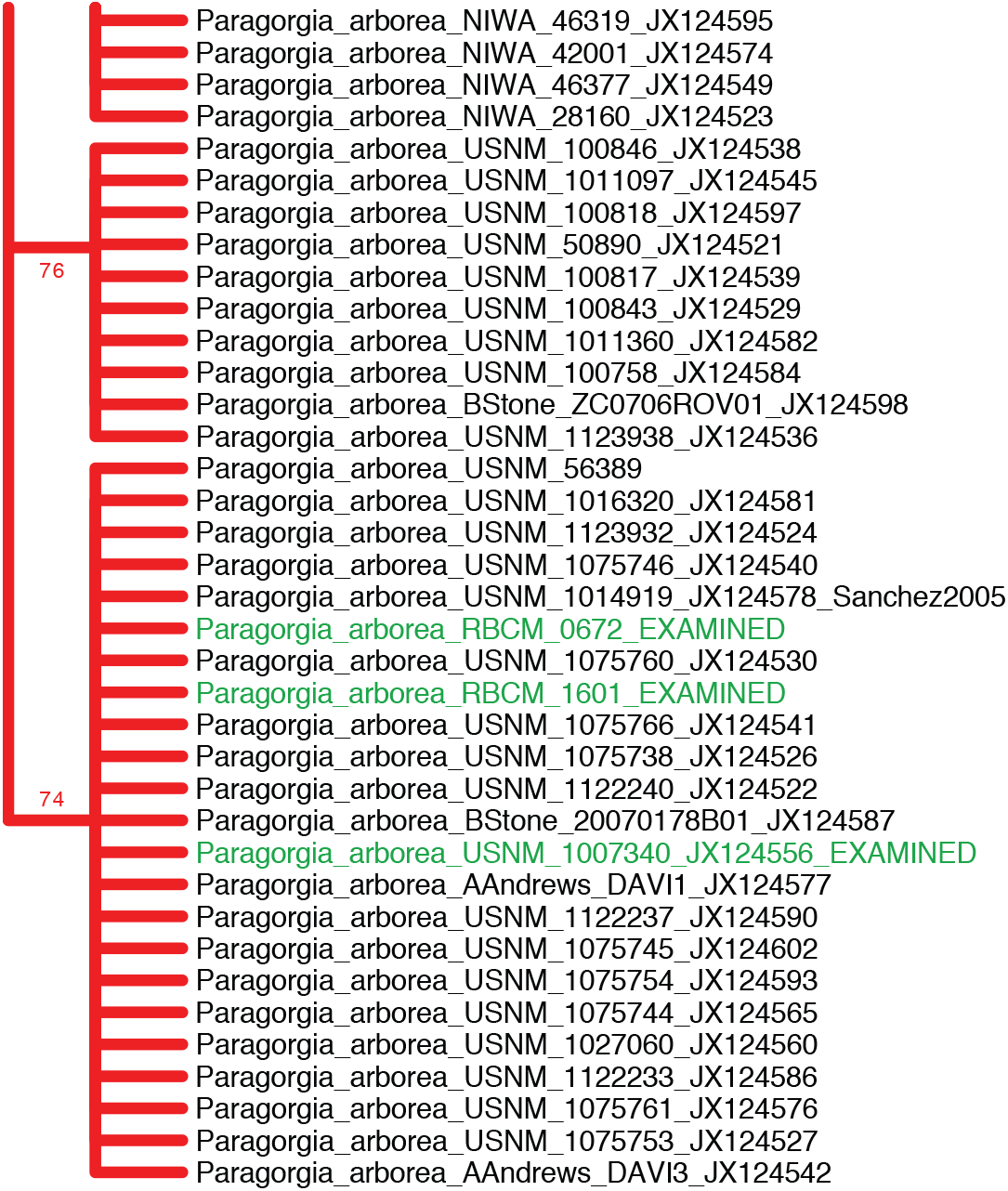
Mitochondrial *mtMutS* bootstrap support consensus tree of all available sequences for the clade AC. Tree inferred from *mtMutS* sequence data from specimens examined in this study, GenBank, and additional specimens. Each family is indicated with a different branch color: red for Paragorgiidae; blue for Coralliidae; and yellow for Alcyoniidae. Outgroups are indicated with black branches. Specimens examined in detail in this study are indicated with green labels. Type specimens are labeled “TYPE”. Specimens examined in Sanchez (2005) are labeled “Sanchez2005”. Tree was created with RAxML using a 50% majority consensus from 500 bootstrap replicates.

**Table S1.**
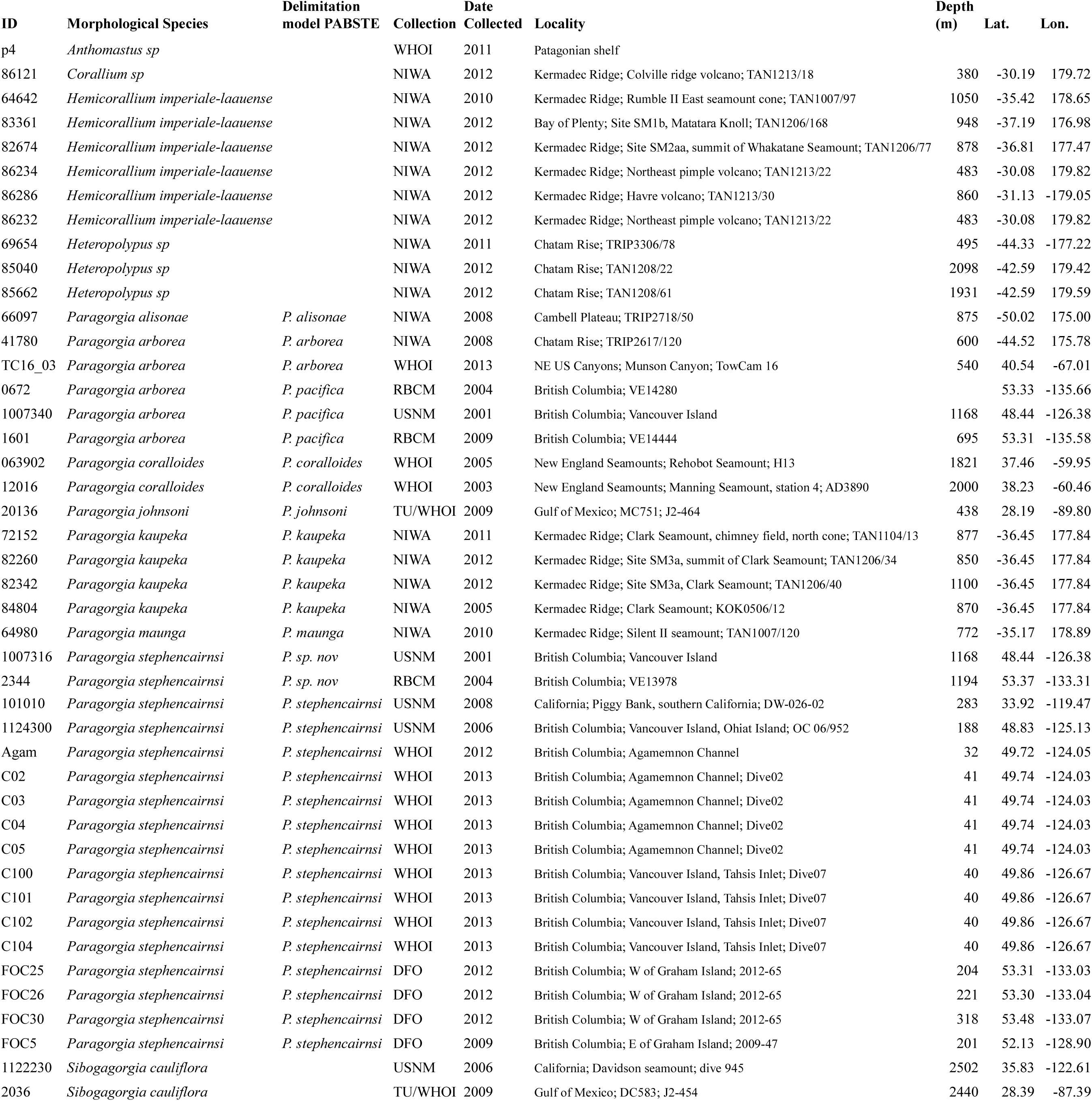
Collection and sequence information for the specimens used in this study.

**Table S2.**
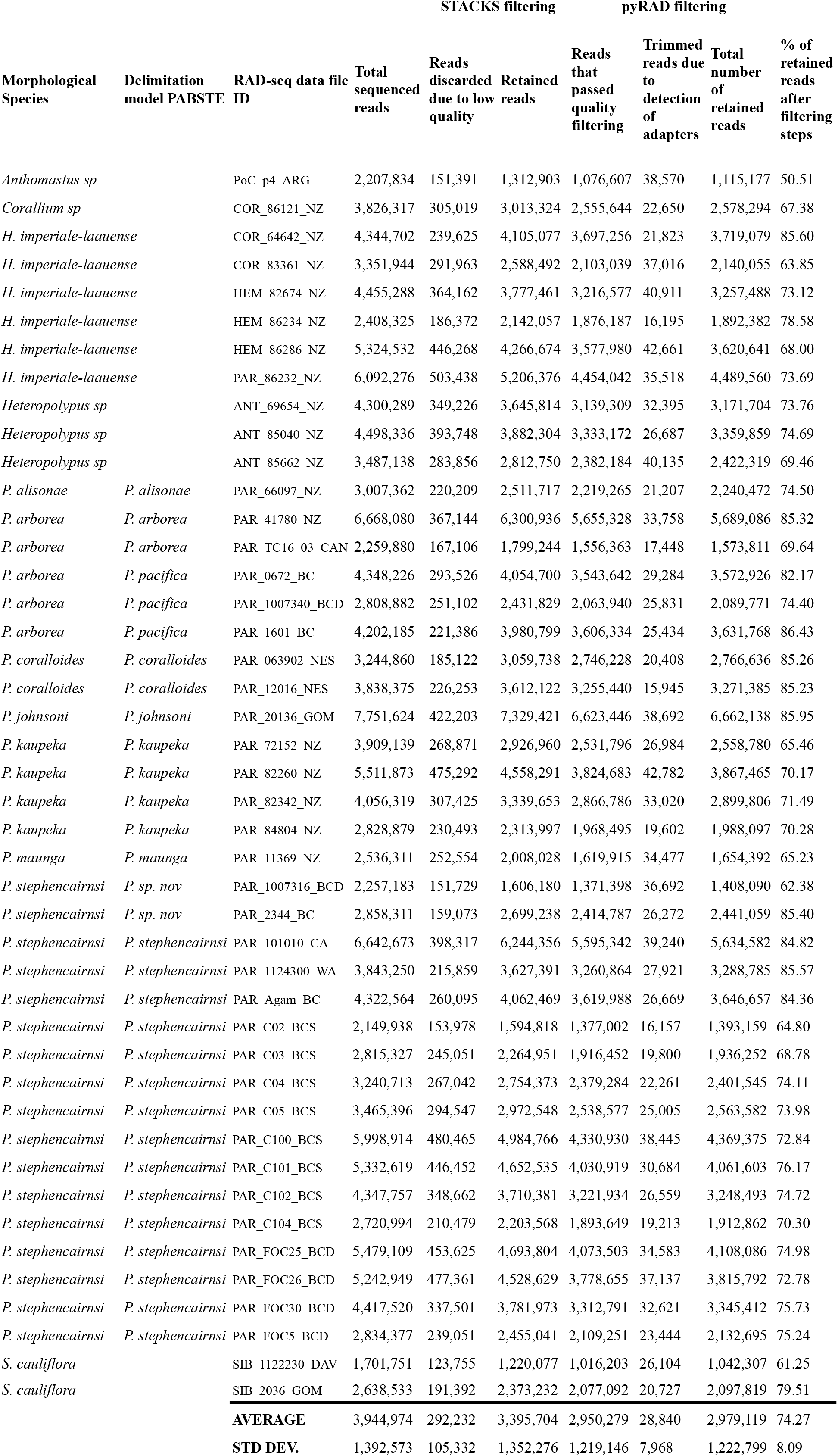
RAD sequencing results and filtering statistics.

**Table S3.**
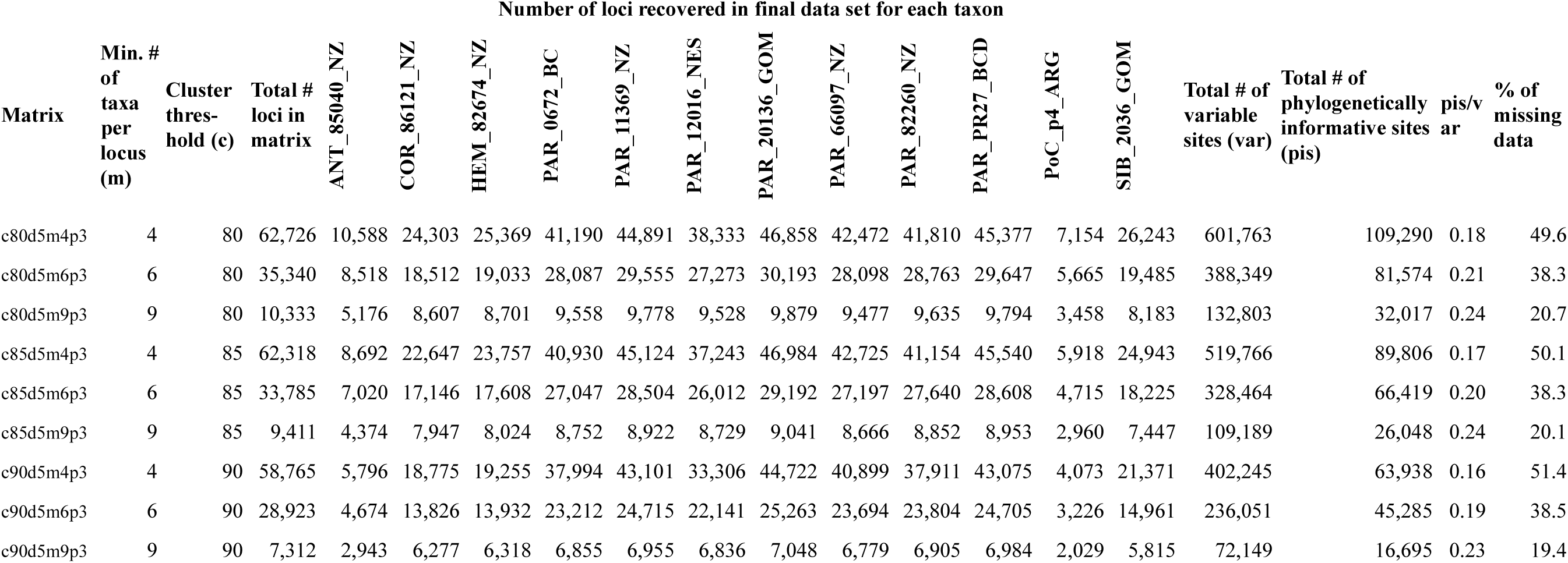
RAD-seq **backbone clustering and matrix statistics.**

**Table S4.**
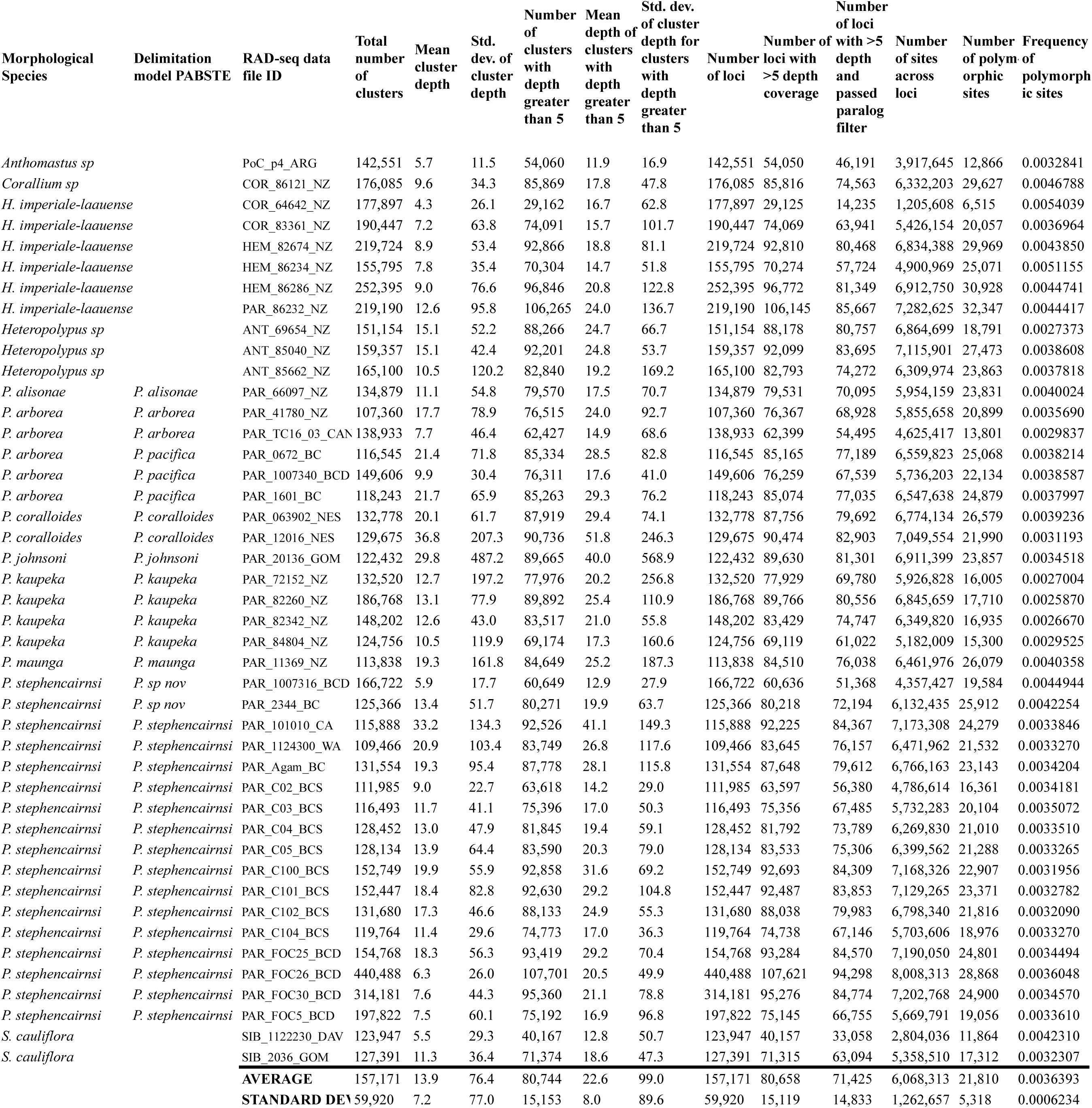
RAD-seq **PHYLO clustering and matrix statistics.**

**Table S5.**
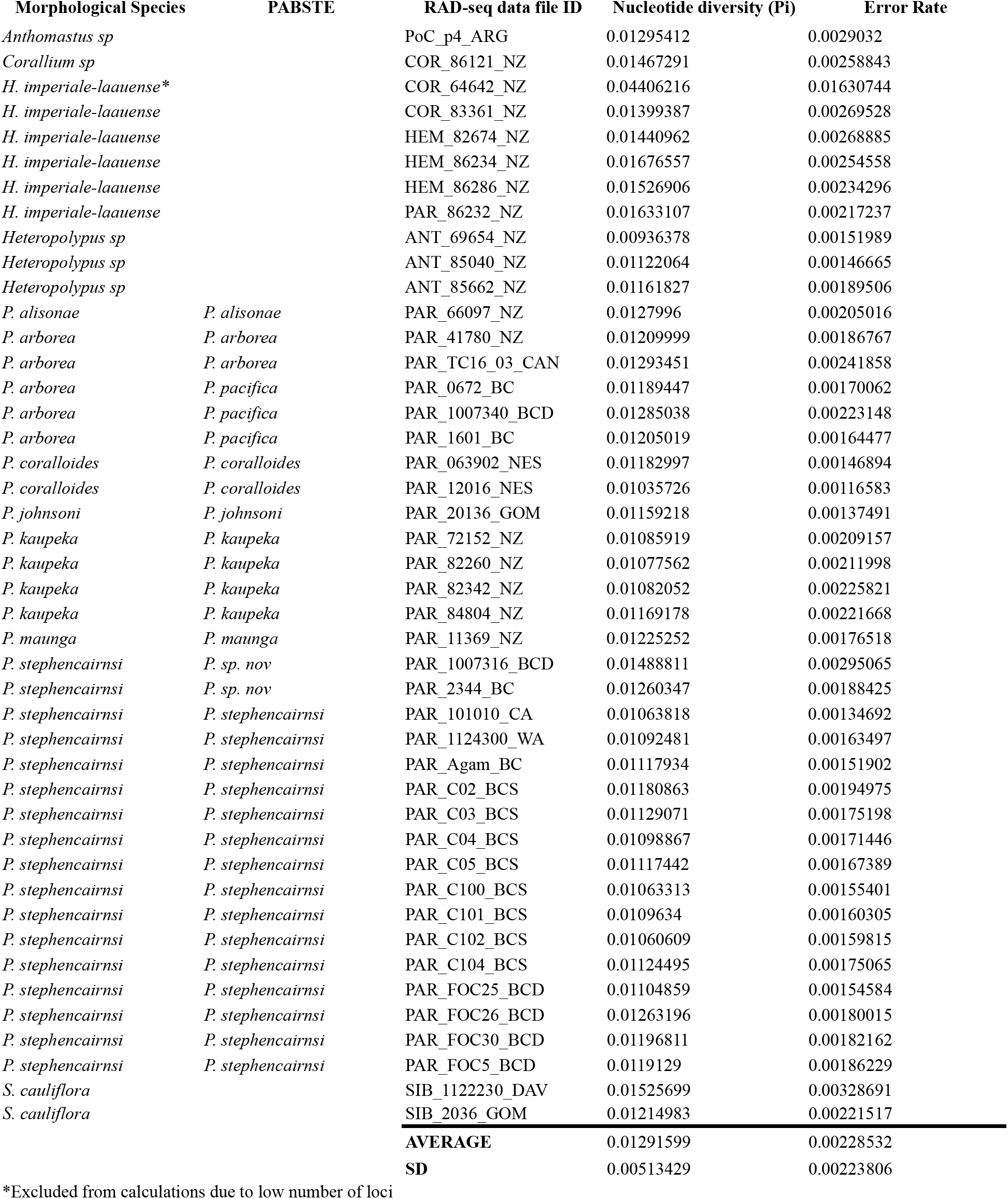
Nucleotide diversity and error rate estimates per specimen based on the **PHYLO matrix**

**Table S6.** Nucleotide diversity and error rate estimates per species based on the **PHYLO matrix**

| Morphological Species | Delimitation model PABSTE | Nucleotide diversity (Pi) |  |  | Per-site sequence error rate (Epsilon) |  |  |
| --- | --- | --- | --- | --- | --- | --- | --- |
|  |  | mean | SD | SE | mean | SD | SE |
| *Anthomastus sp* |  | 0.01295412 |  |  | 0.00290320 |  |  |
| *Corallium sp* |  | 0.01467291 |  |  | 0.00258843 |  |  |
| *H. imperiale-laauense* |  | 0.01935775 | 0.01094005 | 0.00489254 | 0.00447727 | 0.00522005 | 0.00233448 |
| *Heteropolypus sp* |  | 0.01073423 | 0.00120338 | 0.00069477 | 0.00162720 | 0.00023350 | 0.00013481 |
| *P. alisonae* | *P. alisonae* | 0.01279960 |  |  | 0.00205016 |  |  |
| *P. arborea* | *P. arborea* | 0.01245029 | 0.00056582 | 0.00040010 | 0.00193813 | 0.00041487 | 0.00029336 |
| *P. arborea* | *P. pacifica* | 0.01073423 | 0.00120338 | 0.00069477 | 0.00162720 | 0.00023350 | 0.00013481 |
| *P. coralloides* | *P. coralloides* | 0.01245029 | 0.00056582 | 0.00040010 | 0.00193813 | 0.00041487 | 0.00029336 |
| *P. johnsoni* | *P. johnsoni* | 0.01159218 |  |  | 0.00137491 |  |  |
| *P. kaupeka* | *P. kaupeka* | 0.01241323 | 0.00175196 | 0.00087598 | 0.00229768 | 0.00048923 | 0.00024462 |
| *P. maunga* | *P. maunga* | 0.01225252 |  |  | 0.00176518 |  |  |
| *P. stephencairnsi* | *P. sp. nov* | 0.01290646 | 0.00280248 | 0.00198165 | 0.00229281 | 0.00093033 | 0.00065784 |
| *P. stephencairnsi* | *P. stephencairnsi* | 0.01137950 | 0.00066172 | 0.00016543 | 0.00169174 | 0.00016364 | 0.00004091 |
| *S. cauliflora* |  | 0.01370341 | 0.00219709 | 0.00155358 | 0.00275104 | 0.00075783 | 0.00053587 |

**Table S7.**
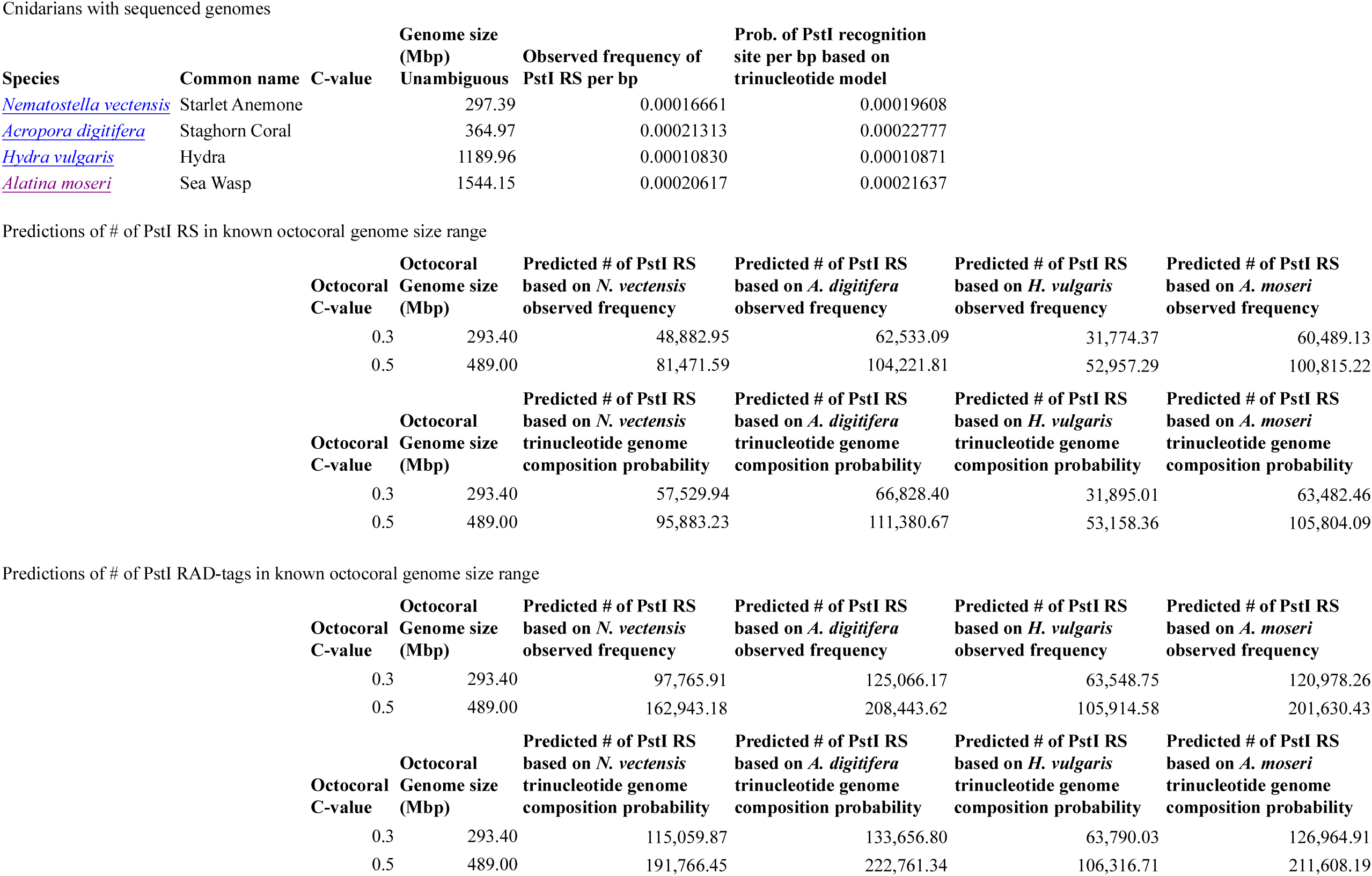
Predictions of # of RAD-tags in octocorals using PstI. Data for *Nematostella vectensis* obtained from the U.S. Joint Genome Institute (JGI-DOE) database. Data for *Acropora digitifera, Hydra vulgaris*, and *Alantina moseri* obtained from the U.S. National Center for Biotechnology Information (NCBI) WGS database. Observed frequency of recognition sequences and calculated probability based on a trinucleotide genome composition model were generated following the methodology described by Herrera et al. (2014). Octocoral genome size ranges were obtained by Luisa Dueñas from gorgoniid octocorals through flow cytomery at the Universidad de los Andes, Bogota, Colombia. Abbreviation: restriction sites (RS).

**Table S8.**
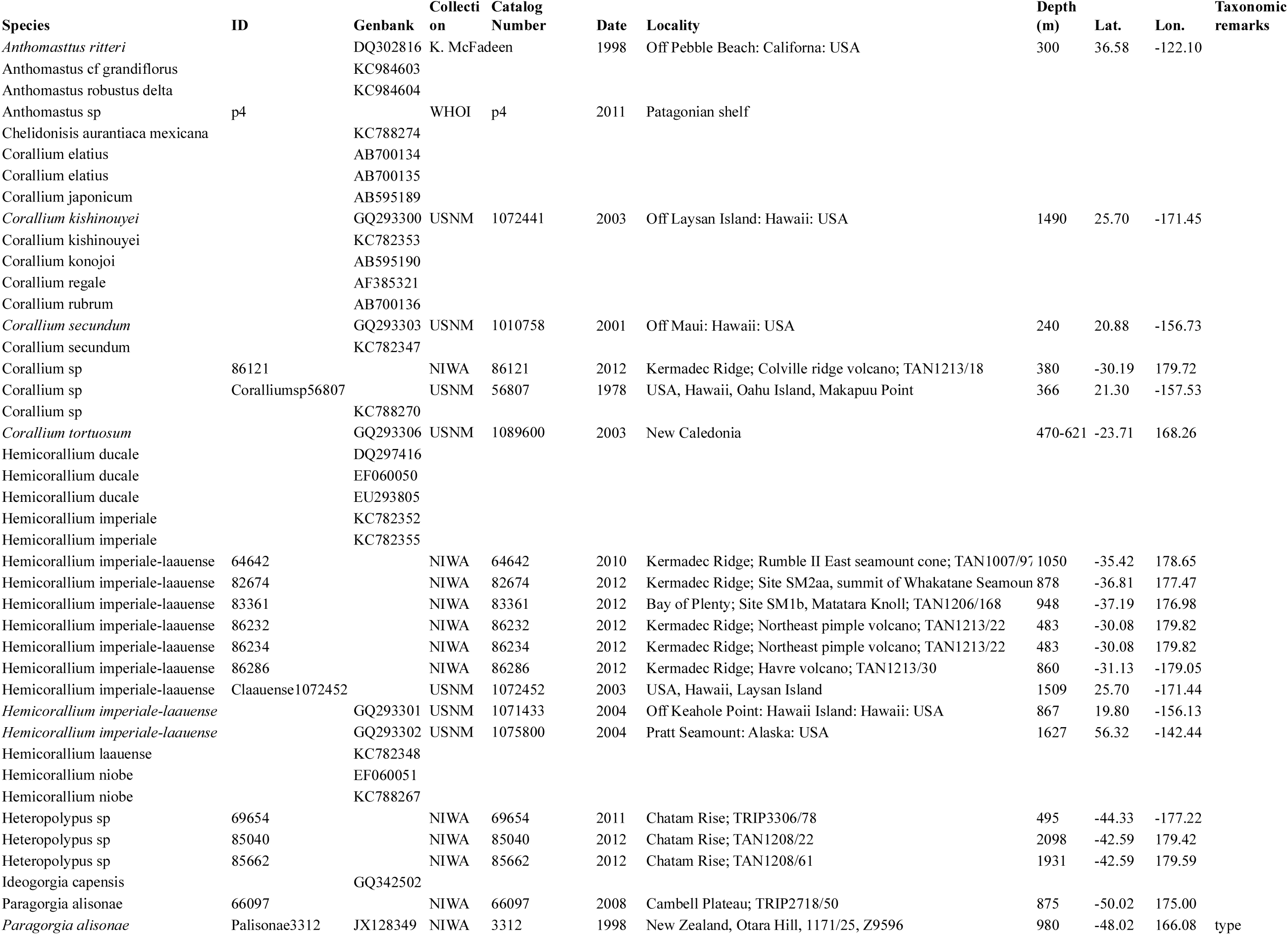
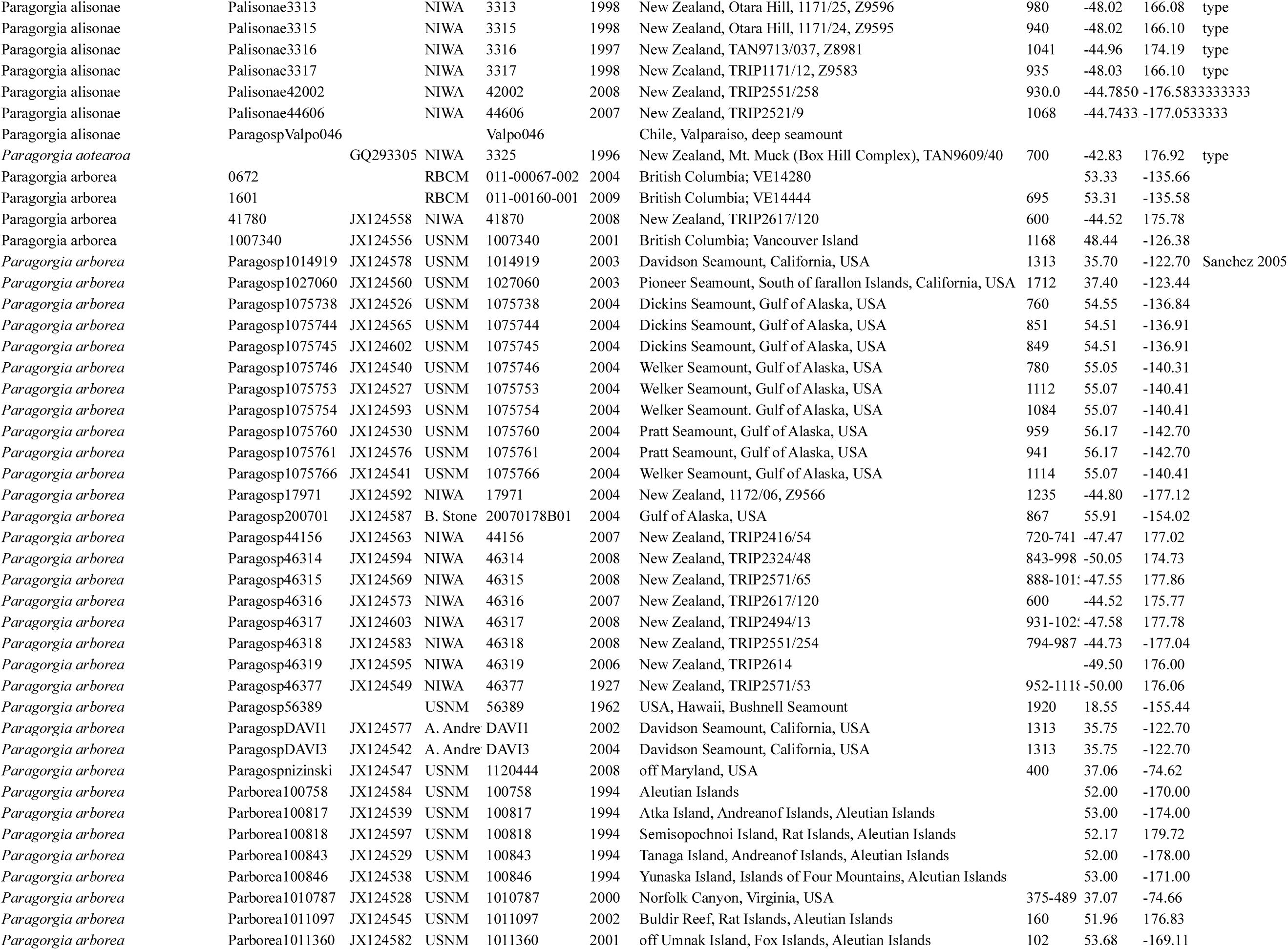
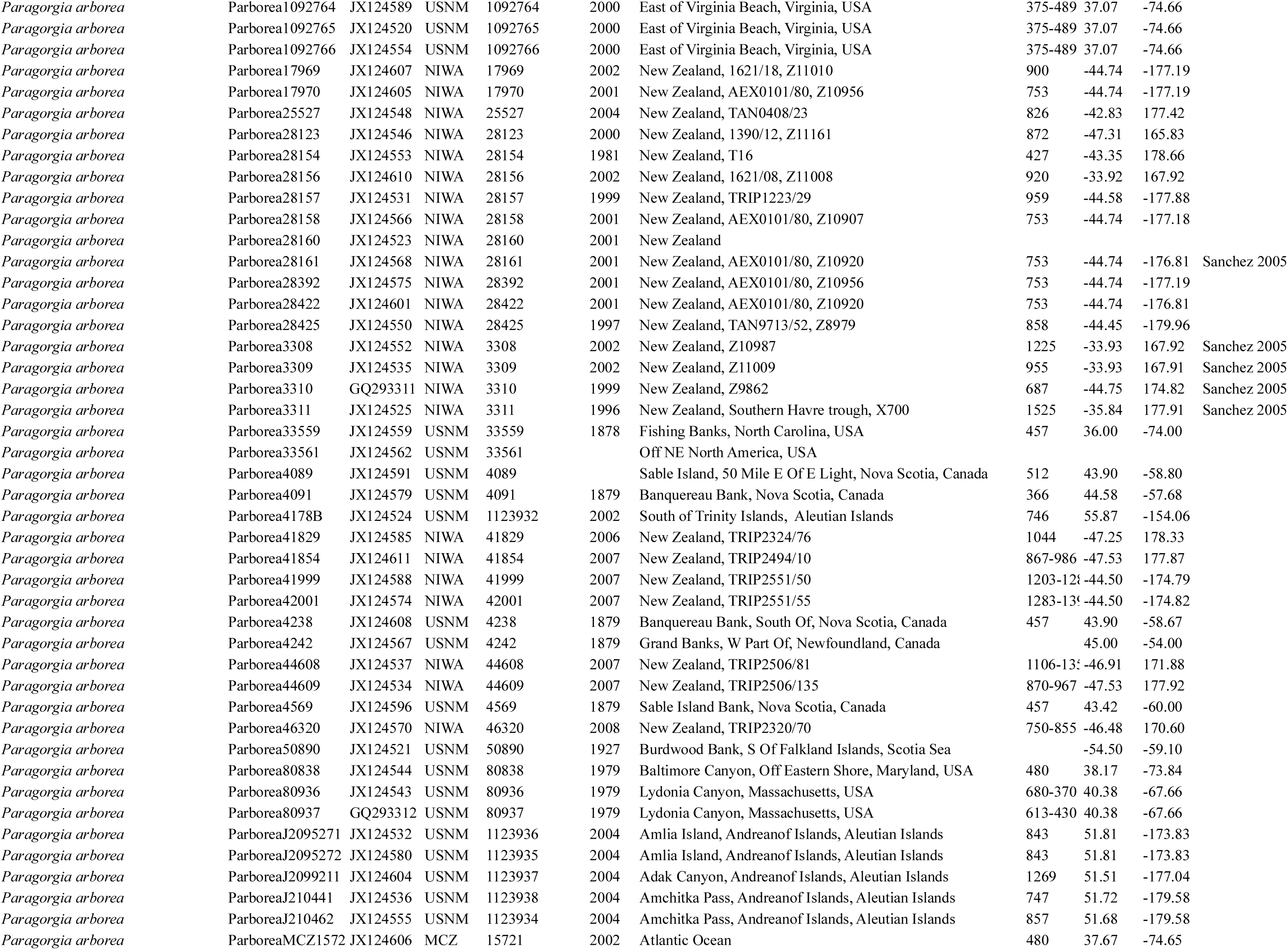
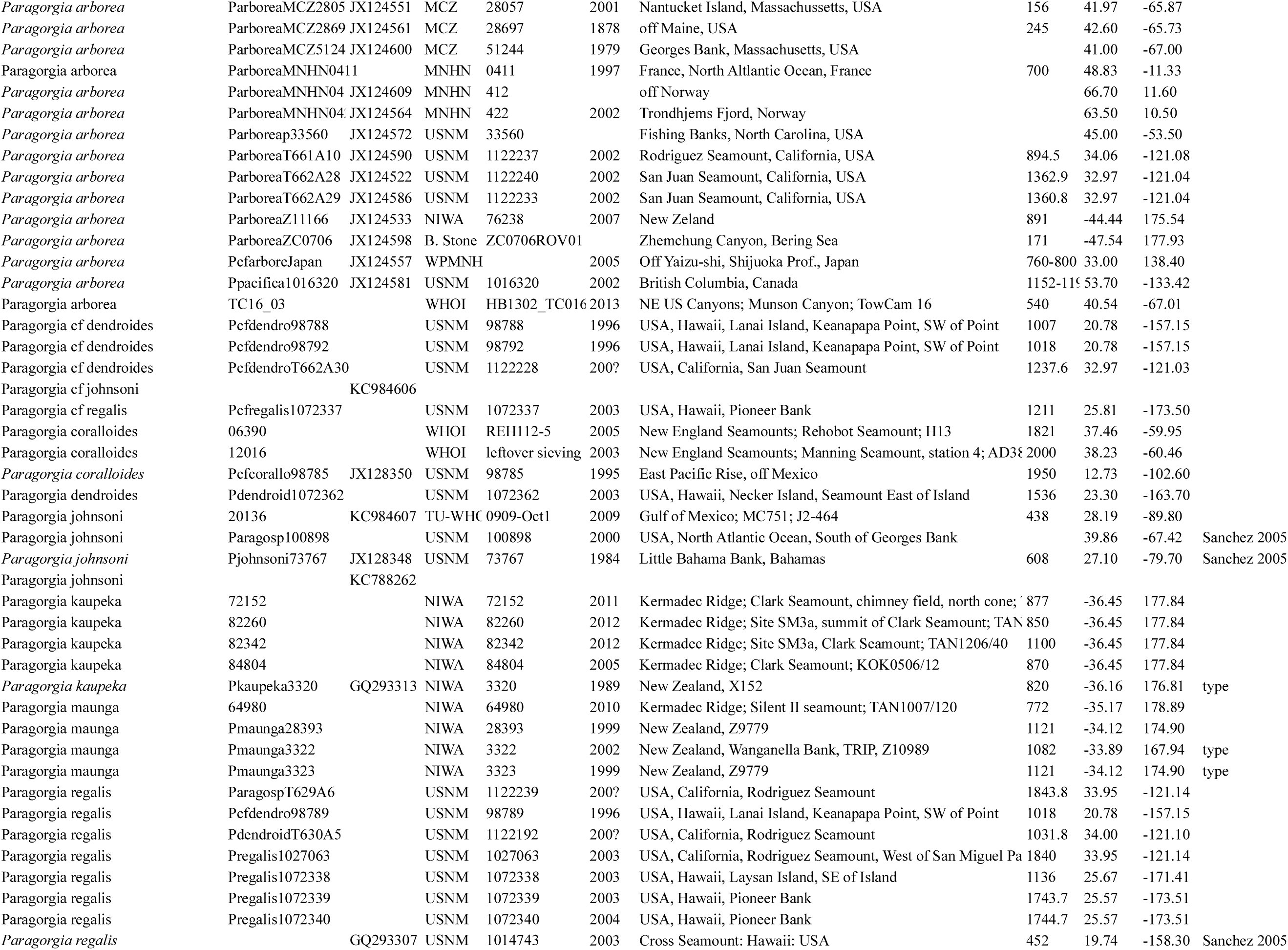
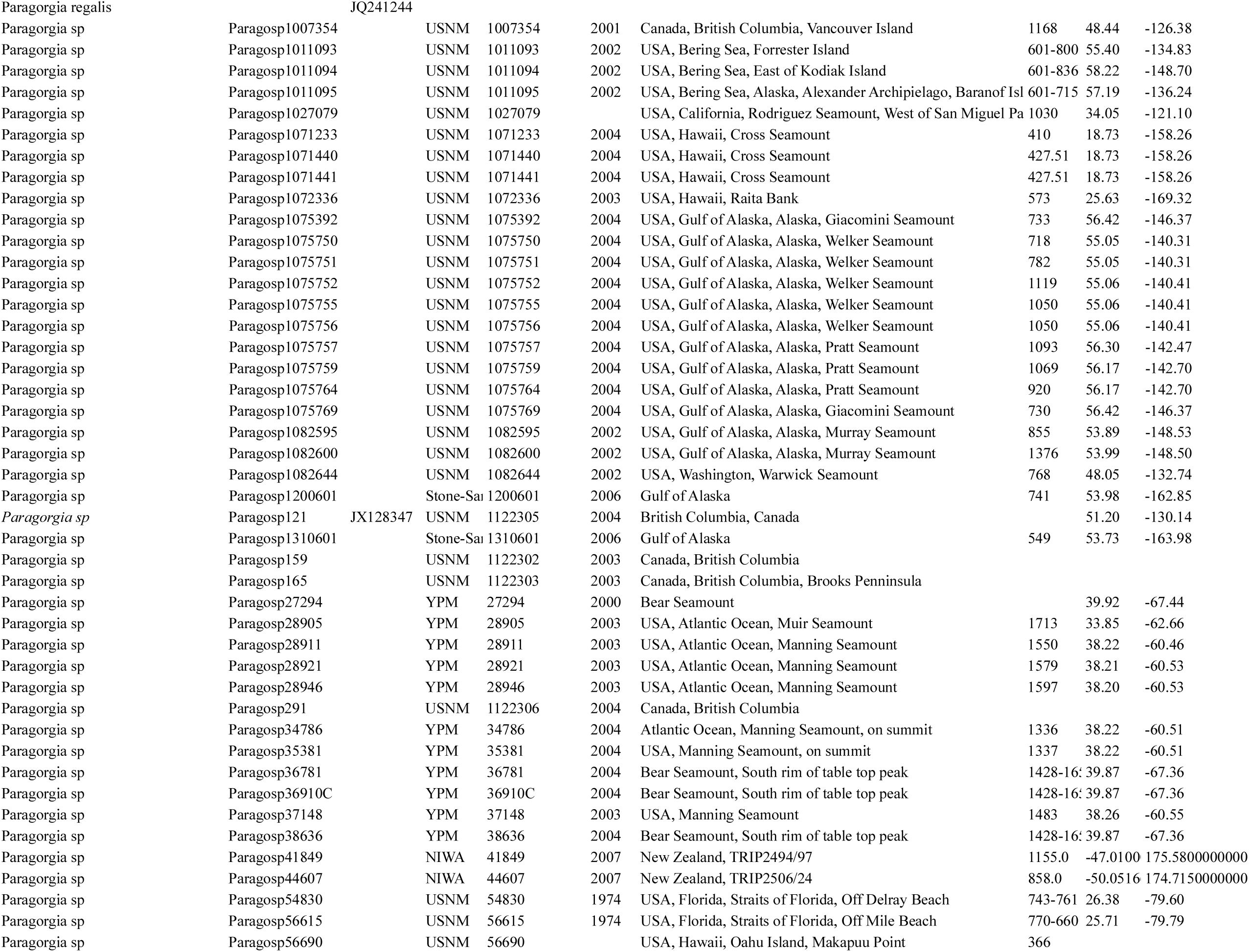
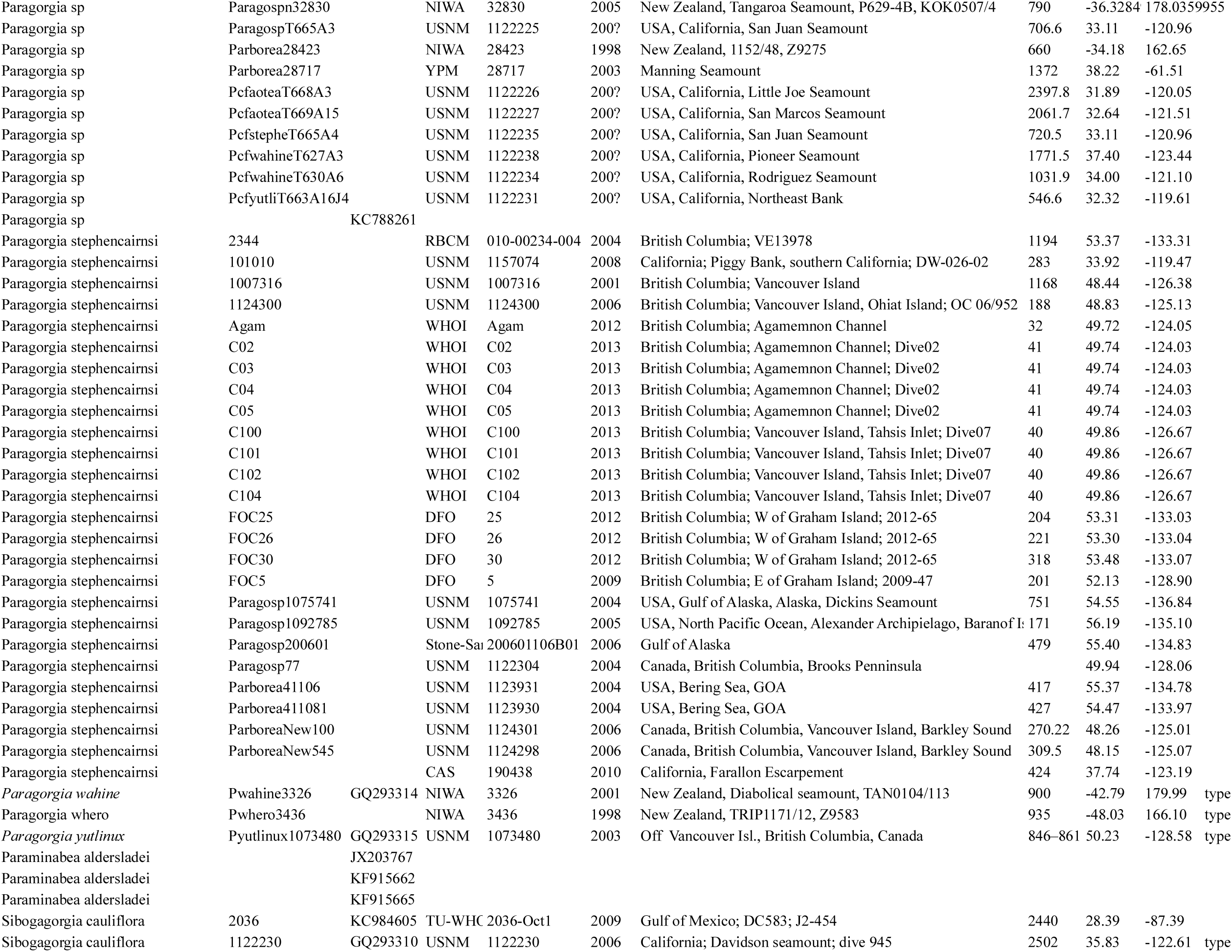
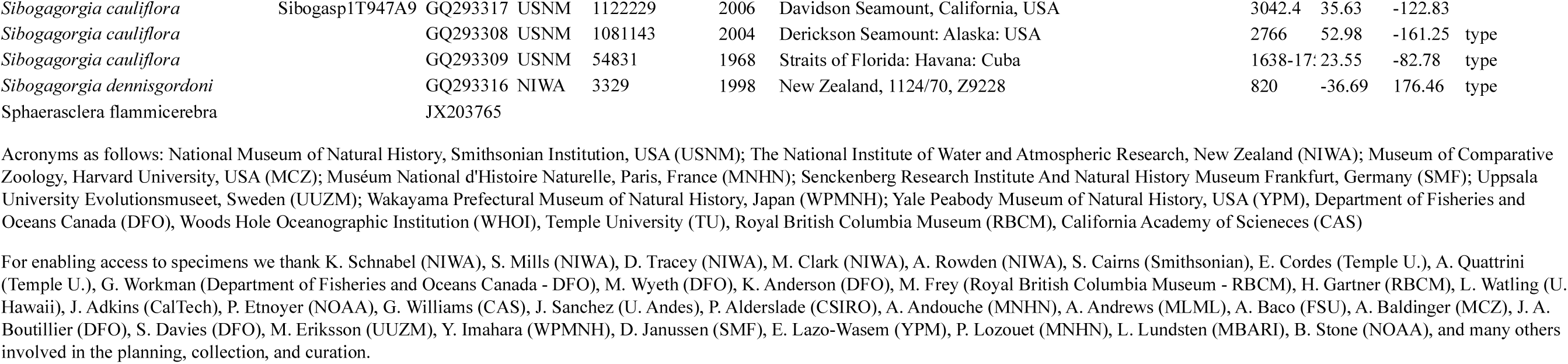
Collection information for all specimens in the clade AC with available mtMutS sequences

